# An ultrastructural map of a spinal sensorimotor circuit reveals the potential of astroglial modulation

**DOI:** 10.1101/2025.03.05.641432

**Authors:** Zachary M. Koh, Ricky Avalos Arceo, Jacob Hammer, Khang Chau, Sarah E.W. Light, Antonio Dolojan, Michał Januszewski, Fabian Svara, Cody J. Smith

**Affiliations:** Department of Biological Sciences and Buchrain, Switzerland; Department of Biological Sciences and The Center for Stem Cells and Regenerative Medicine, Buchrain, Switzerland; Department of Biological Sciences and University of Notre Dame, Notre Dame, IN. Google Research, Buchrain, Switzerland; Department of Biological Sciences and Zürich, Switzerland

## Abstract

Information flow through circuits is dictated by the precise connectivity of neurons and glia. While a single astrocyte can contact many synapses, how glial-synaptic interactions are arranged within a single circuit to impact information flow remains understudied. Here, we use the local spinal sensorimotor circuit in zebrafish as a model to understand how neurons and astroglia are connected in a vertebrate circuit. With semi-automated cellular reconstructions and automated approaches to map all the synaptic connections, we identified the precise synaptic connections of the local sensorimotor circuit, from dorsal root ganglia neurons to spinal interneurons and finally to motor neurons. This revealed a complex network of interneurons that interact in the local sensorimotor circuit. We then mapped the glial processes within tripartite synapses in the circuit. We demonstrate that tripartite synapses are equally distributed across the circuit, supporting the idea that glia can modulate information flow through the circuit at different levels. We show that multiple astroglia, including bona fide astrocytes, contact synapses within a single sensory neuron’s circuit and that each of these astroglia can contact multiple parts of the circuit. This detailed map reveals an extensive network of connected neurons and astroglia that process sensory stimuli in a vertebrate. We then utilized this ultrastructural map to model how synaptic thresholding and glial modulation could alter information flow in circuits. We validated this circuit map with GCaMP6s imaging of dorsal root ganglia, spinal neurons and astroglia. This work provides a foundational resource detailing the ultrastructural organization of neurons and glia in a vertebrate circuit, offering insights in how glia could influence information flow in complex neural networks.

## INTRODUCTION

The precise organization of neuronal circuits is critical for the nervous system. These circuits are established during early embryonic stages. Connectomes of invertebrates like *C. elegans* and recently *Drosophila* have contributed immensely to the understanding of the development and information flow in the nervous system^1–7^. While it is well known that precise connectivity is important, the connectome of even the simplest vertebrate circuits is not complete.

Adding a layer of complexity to vertebrate circuits is the presence of glia. For example, astrocytes in the mammalian brain are predicted to interact with 100K synapses^8^. We know that synapses can be tripartite, involving pre and post-synaptic neurons and astroglia^9,10^. It is also clear that astroglia can modulate synaptic function in multiple ways, including structural support and neurotransmitter modulation. Astrocytic processes that interact with synapses can also exhibit rapid Ca^2+^ microdomains in response to neuronal activity^10–13^. Despite these reports, previous circuit maps have rarely, if ever, included astrocytes. Without an understanding of how neurons and glia are connected in a vertebrate circuit, it is challenging to develop a complete model of how circuits are constructed or how information could flow and be modulated in a vertebrate circuit. This knowledge is foundational and is a necessary resource to understand more advanced topics about circuit development and functionality in vertebrates.

As a model for a simple vertebrate circuit, we focused on the local spinal sensorimotor circuit^14–16^. This sensorimotor circuit is thought to transfer information sequentially from sensory neurons to interneurons to motor neurons, similar to the spinal reflex arc^15,17^. The sensory information from the periphery, however, must be integrated and processed within this circuit before being relayed to the motor neurons. Identifying the complete connectome of a vertebrate sensorimotor circuit is an important step to understand how circuit structure underlies function and supports precise and appropriate motor output in the spinal cord.

We used a serial electron-microscopy data set to reconstruct the glial and neuronal organization of a vertebrate sensorimotor circuit using semi-automated cell reconstructions and automated synapse identification. This work revealed that DRG neurons connect with multiple interneuron populations in the vertebrate spinal cord. The level of complexity in the circuits expands significantly at the interneuron population, with interneurons exhibiting hundreds of synapses. Eventually, interneurons connect with motor neurons. Adding a layer of potential modulation, we demonstrate that astroglial processes are equally distributed across synapses in the circuit. Finally, we validate this connectome resource with functional imaging of GCaMP6s. These findings provide a foundational resource of a vertebrate circuit and demonstrates the potential for complex and multi-layered modulation that includes glia and neurons. Such modulation is critical to ensure animals respond precisely to sensory input.

## RESULTS

To understand the organization of glia within a simple vertebrate circuit, we used the zebrafish sensorimotor spinal circuit as a model^18–22^. We first reconstructed the dorsal root ganglia (DRG) neurons in zebrafish from a serial block-face electron microscopy data set of a 6 dpf spinal cord^23^. This region spanned 74 × 74 × 207 um^3^ (voxel size 9 × 9 × 21 nm^3^)(Fig 1A), which was previously used to skeletonize motor neurons^23^. We employed semi-automated reconstruction approaches to reconstruct neurons in the data set (Fig 1B,C)^24^. While the semi-automated approach was efficient for reconstructions of spinal neurons, a subset of neurons prone to inaccurate segmentation-mergers were manually reconstructed. To determine the exact connectome of the circuit, we also used computational approaches to map all the synapses in the spinal cord section with automated synaptic mapping^24^. Using the reconstructions and synaptic locations, we then systematically reconstructed the vertebrate local spinal cord sensorimotor circuit.

**Figure 1.**
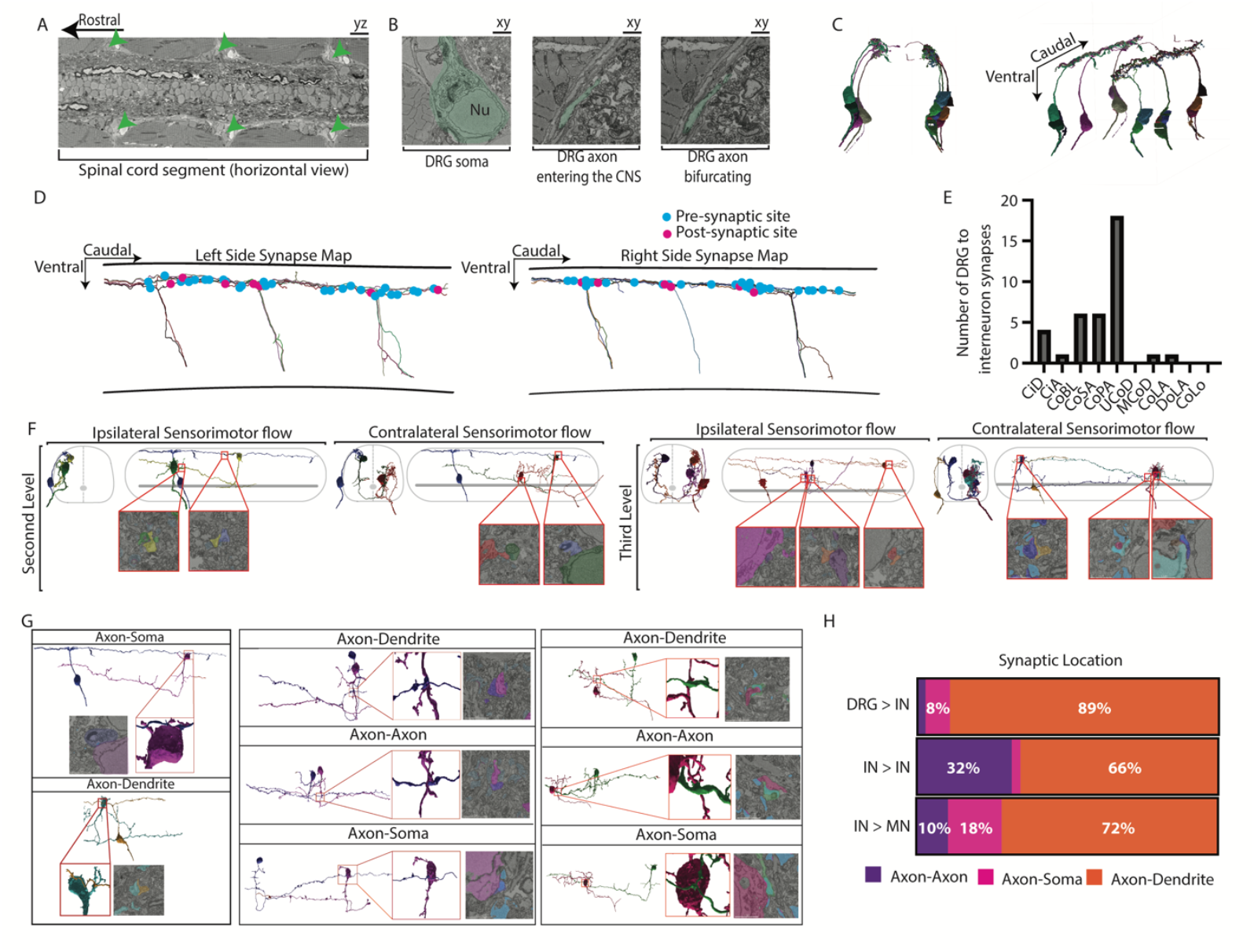
Reconstruction of neurons and synaptic connections within the sensorimotor circuit. A. Electron microscopy (EM) image of the 6 dpf zebrafish spinal tissue that was used to reconstruct the local spinal sensorimotor circuit. Green arrowheads denote the location of the DRG. B. EM images of a DRG neuron (pseudocolored green) at its cell soma (Nu=nucleus), at the PNS/CNS interface, and at its bifurcation location. C. Reconstructions of all the DRG neurons shown in A, demonstrating their precise morphology. D. Reconstruction of the DRG neurons on both sides of the spinal cord overlaid with their synaptic sites. Blue dots denote pre-synaptic sites and pink dots mark post-synaptic sites. E. Quantification of the number of post-synaptic partners of DRG neurons in the tissue section. Note that CoPA synaptic connections are over-represented. F. Reconstructions and EM images of synaptic sites showing that a simple circuit that a DRG, interneuron, and motor neuron could relay information both ipsilaterally and contralaterally. Second level connections indicate a single interneuron connects the DRG with motor neuron, whereas third level requires two interneuron connections. Schematic shows outline of spinal cord and central canal (grey). G. Reconstructions and EM images of synaptic sites at distinct locations of the neurons. H. Quantification of the distribution of synaptic locations at each level of the spinal sensorimotor circuit, showing that most synaptic connections occur between axons and somata at all levels.

In total, 13 DRG neurons were connected into the sensorimotor circuit (Fig 1B,C). The DRG neurons were symmetrically bilateral and extended the length of the tissue section (Fig S1A-E). The DRG were both pre-synaptic and post-synaptic to neurons in the spinal cord and did not display any obvious clustering of synapses (Fig 1D, S1E-I). While synapses are known to occur on secondary branches and were identified elsewhere (Fig S1J-L), we did not detect synaptic sites at secondary branches of DRG neurons. Instead, we found that DRG pre-synaptic sites occurred on the main axonal shaft and thereby represented *en passant* synapses (Fig 1F, S1M). Volume measurements of these synaptic axonal sites revealed local swellings in which synaptic sites were 44% wider than the surrounding axonal shaft (Fig S1N-Q). To identify the specific interneuron subtypes that the DRG neurons connect with, we cross referenced our reconstructions with previous morphological descriptions of interneurons in the zebrafish spinal cord (Fig S2A,B)^19,25–29^. Connections between DRG and CoPA interneurons were over-represented in the circuits (Fig 1E, S2C). Reconstructions of the connected interneurons revealed that *en passant* DRG synapses could form on cell soma and dendritic processes of the interneurons (Fig 1G,H). 196 unique interneurons were connected in these circuits (Fig S2A). The DRG to interneurons connection maps showed information could flow both ipsilaterally and contralaterally through distinct interneurons (Fig 1F). On average, the interneurons exhibited 217.75±78.17 synaptic connections. These diverse interneurons eventually synapsed on 133 motor neurons (Fig 1F,G). Varying subtypes of motor neurons were represented in the map within 3 synaptic connections of the DRG axons (Fig 1F, S2D)^19,23,30,31^. The majority of synapses in the circuit occurred between axons and dendrites but we also detected axon/soma and axon/axon synapses at different levels of the circuit (Fig 1G,H).

To then understand the potential role of glia in a circuit, we extended the map to include glia interactions that could impact information flow through the circuit (Fig 2). In particular, we considered that synapses in the circuit could be tripartite, with pre and post-synaptic neuronal partners in close interaction with astroglial processes^10,32^. The zebrafish spinal cord at 6 days post fertilization (dpf) has astrocytes that are positioned in the synapse rich region where we mapped the sensorimotor circuit^33,34^. To add this additional glial layer, we identified if each synapse in the circuit was contacted by an astroglial process. We defined tripartite synapses as those where an astroglia process was directly adjacent to the synaptic cleft region (Fig 2A). Reconstructions of these astroglial cells demonstrated a robust bushy morphology reminiscent of mammalian and *Drosophila* astrocytes and consistent with morphological characterization from light-microscopy (Fig 2A,B)^10,35–37^. Astroglial processes that interacted with synapses were also present in a subset of astroglia with a radial morphology that extended to the edge of the spinal cord but also had projections that were in the synapse rich region of the spinal cord (Fig 2A,B)^38,39^. While there were astroglial interactions with neuronal processes in non-cleft regions, we did not include such interactions in our analysis. We identified that 23.9% of synaptic connections were tripartite across 3 levels of the sensorimotor circuit.

**Figure 2.**
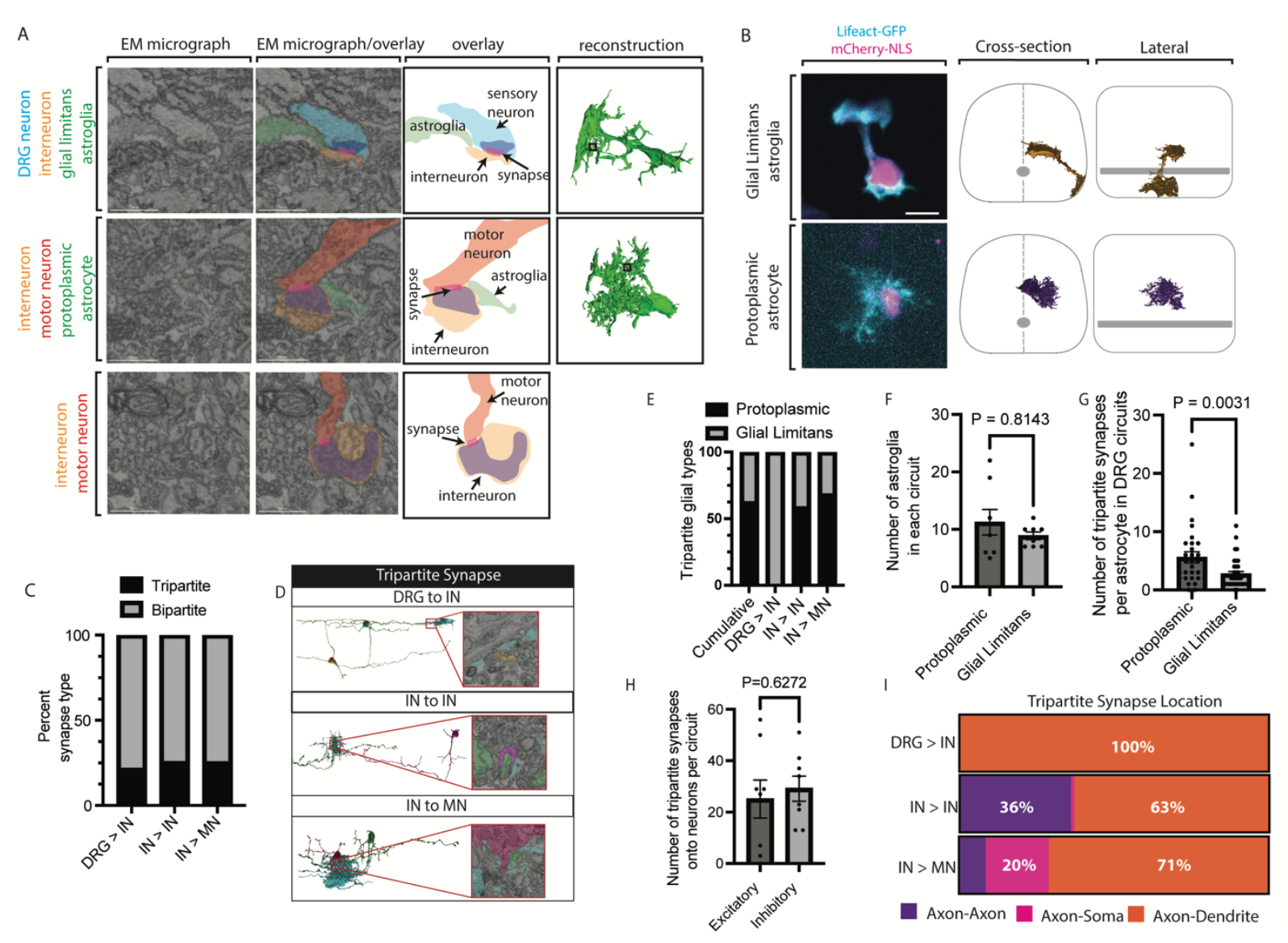
Reconstruction of astroglia in the spinal sensorimotor circuit. A. EM images and cellular reconstructions of synaptic sites within the sensorimotor circuit that do and do not contain astroglia. Note that astroglia can have a radial morphology or a protoplasmic morphology. Pseudocolors correspond with the neuronal subtype. Astroglia are shown in green. B. Confocal images of astrocyte-types in comparison to the EM cellular reconstructions showing distinct astrocyte morphotypes in the spinal cord. Schematic shows outline of spinal cord and central canal (grey). C. Quantification of the percent of synapses that are tripartite vs bipartite between DRG neurons and interneuron (DRG>IN), between interneurons (IN>IN), and between interneurons and motor neuron (IN>MN). D. EM images and cellular reconstructions of DRG/IN, IN/IN, and IN/MN synapses showing tripartite synapses can be at each circuit level. E-G. Quantifications of percent of astroglia subtypes at each level of the circuit (E), number of astroglia subtype seen within individual DRG circuits (F), and number of tripartite synapses per astroglia subtype in the local spinal sensorimotor circuit (G). H. Number of tripartite synapses that are predicted to be excitatory vs inhibitory. I. Distribution of tripartite synapses locations at each level of the circuit. T-test: F,G,H.

Although we know synapses are tripartite, whether specific synaptic connections within a circuit are tripartite is poorly understood. To answer this for the local sensorimotor circuit, we subclassified the astrocyte associated synapses between sensory-interneuron, interneuron-interneuron, and motor-interneuron (Fig 2C,D). These results revealed that tripartite synapses were equally distributed across all levels of the circuit (Fig 2C,D). Both protoplasmic and glial limitans astroglia subtypes interacted with synapses, although sensory neuron to interneuron synapses were exclusively contacted by glial limitans morphotypes (Fig 2E). It is well known that astrocytes have extensive morphology but it is unclear how many astrocytes interact within a given local circuit. To investigate this for the sensorimotor circuit, we asked how many astrocytes were present in each single DRG neuron’s circuit and if single astrocytes form multiple tripartite synapses within the circuit. These results revealed that 20.1±2.5 astrocytes are present within each circuit and a given astrocyte forms 4.06±0.5 tripartite synapses within the circuits (Fig 2F,G). These results support the idea that multiple astrocytes could control synaptic transmission through the sensorimotor circuit or formation of it. To identify if tripartite synapses were positioned in specific areas of a single circuit, we subcategorized the tripartite synapses in various ways according to their neuronal partners. We first asked if tripartite synapses were biased toward pre-synaptic partners that would be expected to be excitatory vs inhibitory (Fig 2H). We assumed Dale’s law – each neuron has only one neurotransmitter profile^40,41^. Tripartite synapses were present at 46% of predicted excitatory vs 54% of inhibitory neurons, suggesting equal distribution across neuron types. Finally, we determined tripartite synapses were located at synaptic connections between axon-dendrite, axon-soma or dendrite-dendrite (Fig 2I). Together these results reveal the precise and extensive network of glial interactions within a single circuit that could impact the processing of sensory information.

Now that the glial and neuronal organization was mapped, we compiled it to understand the potential information flow. We represented the circuit with: reconstructions at different levels, a heat map of connected neurons, and a Sankey plot that showed information flow based on synaptic connections (Fig 3A-C). Level 1 connections were interactions directly between DRG neurons and their partners (Fig 3A,C). The second level represents the interconnectivity of interneuron connection (Fig 3A-C). Finally, the third level exhibited the extensive connections that ended at motor neurons (Fig 3A-C). Even looking at a single DRG neuron’s circuit reconstructions demonstrated the potential for complicated flow of information. In the representative DRG neuron, the neuron is connected to two excitatory neurons, CoPA and CiA and an inhibitory neuron CoSA (Fig 3A,C). Examining the second level connections identifies that information from a single DRG neuron can travel to a motor neuron on the ipsilateral side through the CiA neuron (Fig 3A). It also revealed that the CoSA neuron could relay that information to the contralateral side of the spinal cord (Fig 3A). The second level of connections demonstrated a network of interneuron to interneuron connections that increased the complexity of the circuit (Fig 3A). Finally, the third level of connections demonstrated that information could travel both ipsilaterally or contralaterally to eventually impact motor neurons (Fig 3A). For even a single DRG neuron, the circuit is complex, connected through 155 synapses (Fig 3A-D). For example, the motor neuron on the ipsilateral side can be excited by an interneuron. The activity of the motor neuron on the contralateral side can be changed as well, either by predicted excitatory interneurons that are commissural or through two inhibitory neurons in succession (Fig 3C,D). The number of times a cell is synaptically connected in a circuit could indicate its influence in that circuit. To probe this concept and determine how many cells and synapses per cell are present within a single DRG neuron’s circuit, we aggregated those numbers into a heat map (Fig 3B). The heat map demonstrates that while many cells are only connected in the circuit through one synapse, others make multiple synaptic connections (Fig 3B). Aggregating the classes of neurons in a Sankey plot provided an understanding of how neuronal information could flow through the local circuit. For example, the local circuit relays information directly to motor neurons within two synaptic connections but also expands significantly through the connections of CiA interneurons (Fig 3C, S3A). Lastly, using previous studies that identified the interneurons class with excitatory vs inhibitory^19,25–29^, we mapped the predicted excitatory vs inhibitory neurons that are present in a single DRG neuron’s circuit (Fig 3D). This demonstrated that excitatory information could flow quickly to motor neurons and further refined the understanding of how information flows within the local spinal circuit.

**Figure 3.**
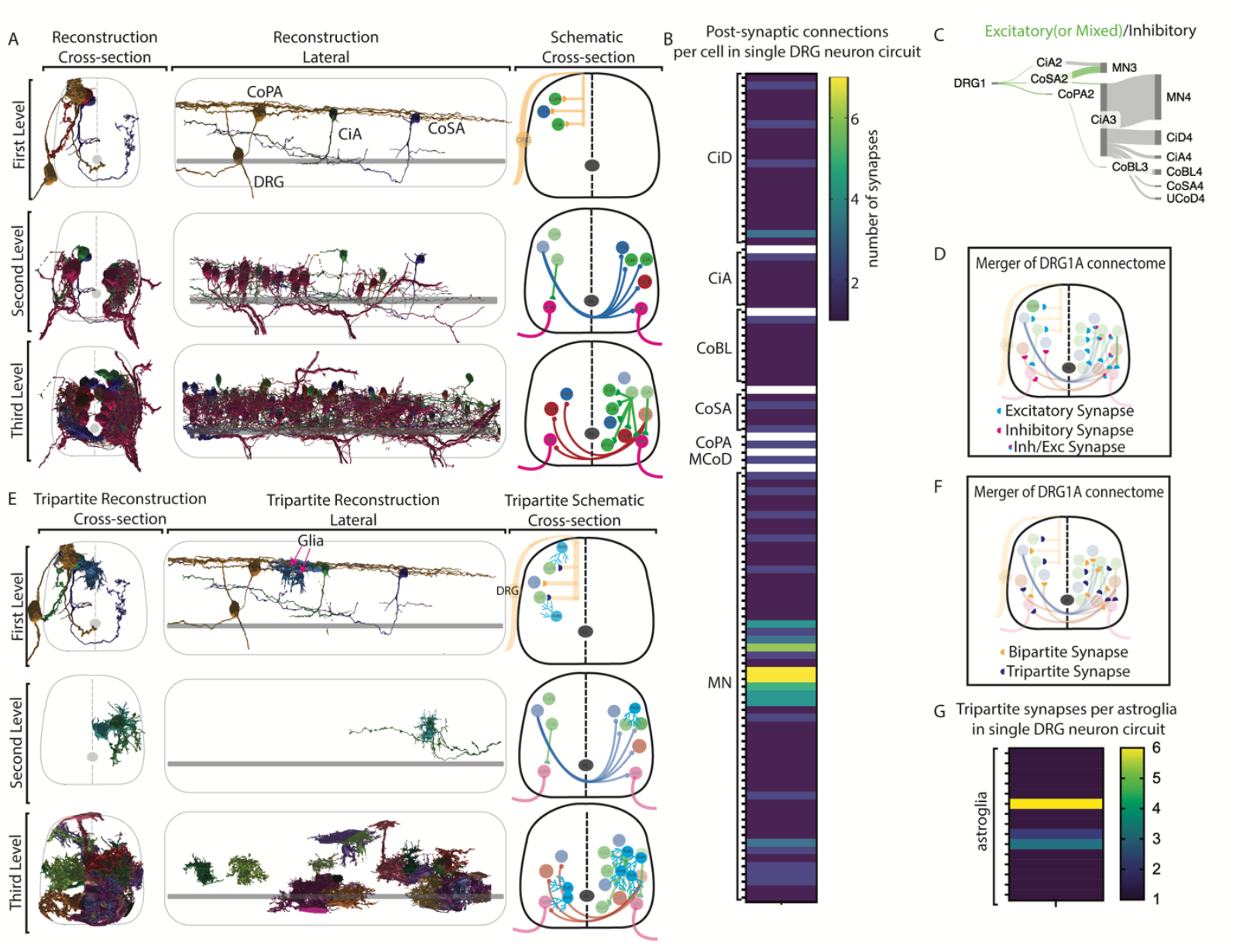
Neuronal and astroglia organization in single sensorimotor circuits. A. Cellular reconstructions of neurons in the local spinal sensorimotor circuit from a single DRG neuron’s circuit. Schematic representation of the neurons is color coded based on the excitatory (green), inhibitory (red), or mixed (blue) identity of the neurons, DRG neurons (yellow), and motor neurons (pink). Schematic shows outline of spinal cord and central canal (grey). B. Heat map showing the number of synapses for each cell in a single DRG neuron’s circuit. Each row represents an individual neuron of the bracketed identity. C. Schematic showing potential information flow demonstrating each of the classes of neurons and how that could be impacted when incorporating excitatory or inhibitory neurons. The size of the bar represents the number of synapses that connect the classes of neurons. D. Schematic of a single DRG neuron’s local circuit demonstrating the predicted excitatory (blue), inhibitory (magenta) and mixed connections (blue/magenta). E. Cellular reconstructions of neurons and astroglia in the circuit depicted in A. All astroglia (sky blue) that are associated with the synapses in that circuit are included. F. Schematic showing tripartite (purple) vs bipartite synapses (orange) within a single DRG neuron’s circuit that is depicted in A. G. Heatmap of the number of synapses that each astroglia interacts with in the DRG neuron circuit depicted in A. Each row represents an individual astroglia of the bracketed identity.

To ask how astrocytes could contribute to the complexity of the circuit, we added the astrocytic interactions onto the circuit of a single DRG neuron (Fig 3E, S3B). These results revealed where astrocytes could modulate the circuit. In just the 3 levels, astrocytic processes contacted multiple synapses. In particular, we revealed that astrocytes could modulate the first level of synaptic connections between the DRG neurons and the interneurons (Fig 3E). To represent the control of astrocytes on a single DRG neurons circuit, we collapsed the synapses onto a single map (Fig 3F). These results revealed a network of synapses that would be expected to be excitatory and inhibitory neurons (Fig 3D). The collapsed circuit demonstrates a precise map of tripartite synapses vs bipartite synapses (Fig 3F). It is possible that astrocytic modulation could be controlled by a collection of astroglia or by a single highly-connected astroglia. To explore this, we created a heat map of how many times each astrocyte participated in a tripartite synapse within a single DRG neuron circuit (Fig 3G). Although a portion of astroglia only form one tripartite synapse in the circuit, at least 3 astroglia had the potential to modulate multiple synapses within the circuit (Fig 3G). Incorporating astrocytes into the map of the sensorimotor circuit demonstrates a complex circuit that could be modulated by both neurons and glia at multiple layers.

Spinal somatosensory responses have spatial acuity but interneurons also extend through multiple hemisegments, suggesting that spatial acuity might be in part coded by interconnected circuits. To next explore the extent of interconnectivity, we compiled the maps of all the DRG neurons in the tissue segment (Fig 4). To understand the scale of information flow at each level, we created a Sankey chart where the number of synapses is collapsed at each level (Fig 4A, Fig S4A). These results demonstrated that the synaptic connections expanded significantly at the interneuron level (Fig 4A). This potential information flow funneled into the motor neurons. Layering glial cells onto this information flow map demonstrated that glia had potential to modulate the circuit at all phases equally (Fig 4A). To next understand the directional flow of potential information, we generated a map where individual synapses of DRG neurons are mapped onto interneuron and motor neuronal subtypes (Fig S4A). These results demonstrated DRG neurons’ synapse onto most interneuron subtypes (Fig S4A). Even within the 3 segments of the spinal cord that we mapped, there was massive interconnection between DRG circuits. For example, synapses within the integrated sensorimotor circuits existed between CiA, CoSA, CoPA, CoBL, CiD, MCoD, UcoD, COLA interneurons and motor neurons (Fig S4A).

**Figure 4.**
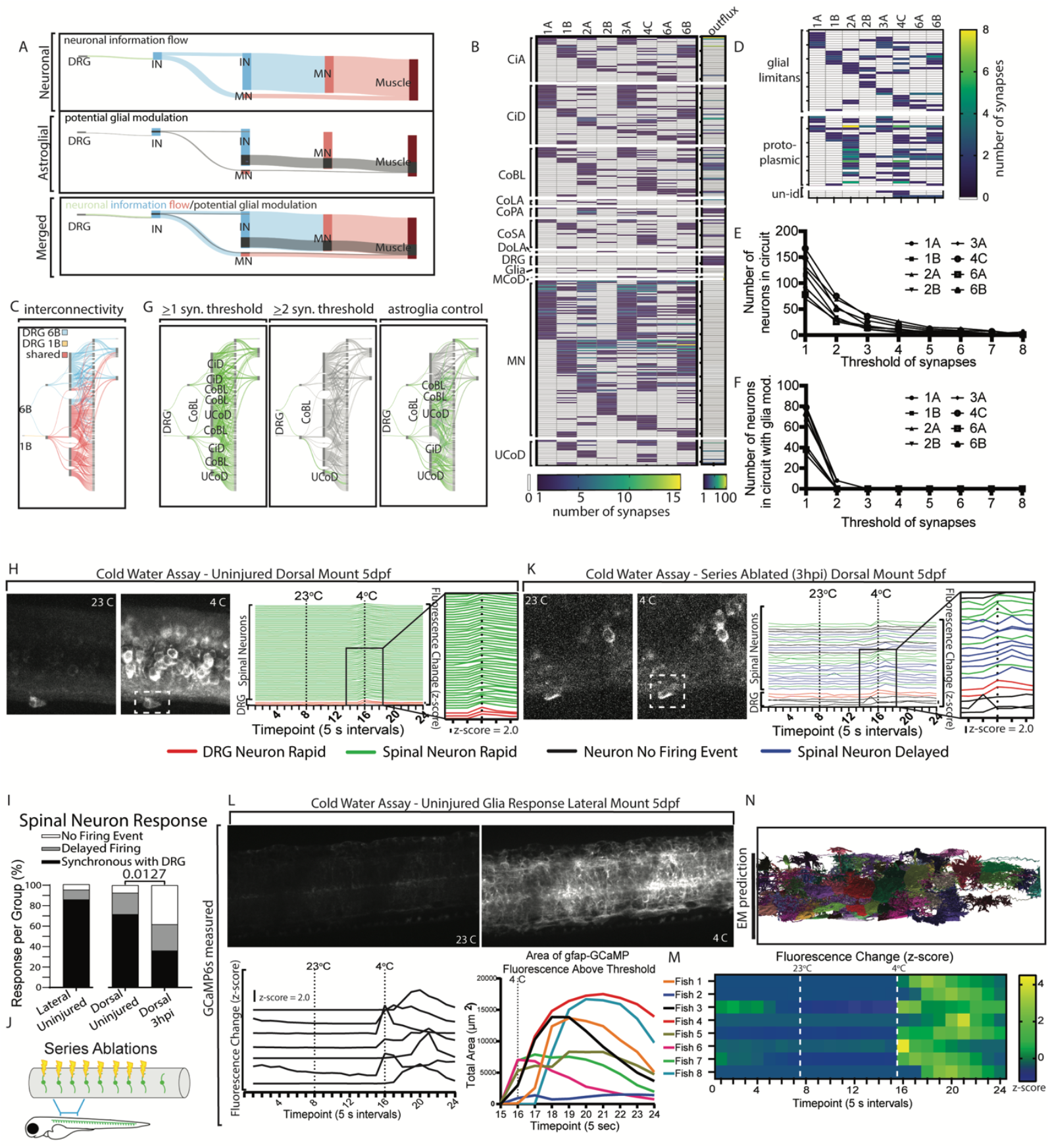
Spinal sensorimotor circuits are interconnected. A. Sankey plot of the DRG, interneuron (IN), and motor neuron (MN) connections aggregated from all the DRG in the tissue section to show where information can expand in the circuit. The height of the bar represents the number of synaptic connections at that layer of the circuit. B. Heatmap of every neuron that is connected within the local spinal sensorimotor circuit. Colors denote the number of synapses on each given neuron within each circuit. Columns are organized to demonstrate each DRG neuron’s circuit. Classes of neurons are clustered together. Outflux column shows the cumulative amount of post-synaptic connections a cell has in all the circuits C. Sankey plot of DRG 1B and 6B with the lines representing information flow are colored based on whether they are unique to 1B (orange), 6B (blue), or shared (purple) to demonstrate how interconnected each DRG neuron’s circuit is. D. Heatmap of every astroglia that is connected within the local spinal sensorimotor circuit. Colors denote the number of synapses that each astroglia interacts with in each single DRG neuron’s circuit. E. Quantification of the neurons in each DRG neuron’s circuit if information flow was controlled by the number of synapses required to induce neuronal activity. Note that as the threshold of synapses increases, the number of neurons in a circuit decreases. F. Quantification of the neurons in each DRG neuron’s circuit if flow of information was controlled by astroglia. G. Sankey plot of potential information flow if the number of synaptic connections on a given neuron controls the flow. Represented are the interconnected circuits of 1B and 6B depicted in C. >1 and >2 demonstrates the drastic change in information flow when synapse thresholding is applied. Also depicted is the information flow if astroglia could modulate the circuits of 1B and 6B. Astroglia control plot assumes synaptic thresholding is >1. H. Confocal images and quantification of DRG and spinal neurons when *Tg(neurod1:Gal4); Tg(UAS:GCaMP6s)* animals are exposed to 23°C and 4°C water. Quantifications are represented as z-score changes of intensity. DRG neurons with a rapid change in z-score >2.0 are colored red, spinal neurons with rapid changes are colored green, spinal neurons with delayed changes are blue, and neurons that do not change (z-score>2.0) are black. Note that exposure to 4°C water causes DRG and spinal neurons to rapidly increase GCaMP6s intensity. I. Quantifications for the percent of neurons that have each z-score change characteristic. Bars represent lateral mounted, dorsal mounted, and dorsal mounted injured animals. P=0.0127, Fisher’s exact test between dorsal uninjured vs injured conditions. J. Schematic of the serial axotomy paradigm where 8 consecutive nerves that connect the DRG with the spinal cord are severed. K. Confocal images and quantifications of DRG and spinal neurons of *Tg(neurod1:Gal4); Tg(UAS:GCaMP6s* animals that previously underwent serial axotomy. Note that DRG neurons still show z-score >2.0 after 4°C but fewer spinal neurons show rapid z-score changes >2.0. L. Confocal images and quantification of the fluorescent intensity and area when *Tg(gfap:GCaMP6s-CAAX)* animals are exposed to 23°C and 4°C water. Note that astroglia exhibit both changes in GCaMP6s fluorescence intensity and area after exposure to 4°C water. M. Z-score calculations of 8 *Tg(gfap:GCaMP6s-CAAX)* animals exposed to 23°C and 4°C water showing that the change in fluorescence is specific to 4°C exposure. N. Cellular reconstructions of all astroglia that interact with synapses in the aggregate DRG neuron’s circuits. Fisher’s Exact test: I

Finally, to understand how individual neurons were represented in the integrated map, we generated heat maps where each neuron in the collective circuits was colored based on how frequently it was represented within the circuits (Fig 4B). The heat map revealed that individual DRG circuits contain both unique synaptic connections for that circuit and shared synaptic connections with other DRG circuits (Fig 4B). An example of this is DRG neurons 1B and 6B which are on opposite sides of the spinal cord but share a common CoBL neuron (Fig 4B,C, S4B). While there is convergence onto single neurons, the DRG neuron circuits also display circuit divergence. This is prominent with CiA and CiD neurons, which are typically exclusive to a single DRG neuron’s circuit (Fig 4B). This concept is highlighted by again contrasting DRG neurons 1B and 6B, with 6B connecting to 73 unique connections between neurons compared to 1B (Fig 4C). To determine the potential of glia to modulate this interconnectivity, we asked how glia are positioned within these interconnected circuits by performing a similar analysis for glia. This showed distinct astroglia that had potential to modulate multiple synapses within a single circuit and between distinct local circuits (Fig 4D). Interestingly, protoplasmic astrocytes were more interconnected than glial limitans astroglia (Fig S4C). Together, these results demonstrate a complex sensorimotor circuit that has both potential for convergent and divergent information flow through spinal neurons that could be modulated by glia.

With the massive interconnectivity of the neurons in local circuits, it seemed possible that the simple connectivity of neurons could not provide acuity in sensory responses. To explore how sensory acuity could be coded in the spinal cord, we considered the hypothesis that the number of synaptic connections for a given neuron could refine the number of neurons involved in a circuit^6,7,42,43^. To test this idea, we determined how many neurons were in each circuit if the threshold for firing a neuron was based on the number of synaptic inputs. While the circuit was complex if an input threshold of one synapse was required, connecting 116+/-34.6 neurons, thresholding the system to 3 synapses reduced the number of neurons in the circuit to 21.8+/-11.1 (18.4+/-6.6% of the total cells) (Fig 4E). To determine how glia could function in this thresholding, we created similar calculations by subtracting the number of synaptic connections that contained glia. Thresholding with glia reduced the circuits on average from 116+/-34.6 to 60+/-6.9 neurons without any additional synaptic thresholding (Fig 4F), demonstrating the potential control that glia could have on circuit complexity and significantly reducing the scale of thresholding required to simplify a circuit.

While the number of neurons in a circuit provides an understanding of circuit complexity, it does not incorporate the directional flow of information through circuit levels. To ask how the information flow could be altered from thresholding of synaptic number, we generated flow charts of every neuron in two interconnected DRG circuits (1B and 6B)(Fig 4G). Theoretically, circuit information could flow through 217 connections between neurons if only 1 synapse is required (Fig 4G). Thresholding the synapses to >2 downstream of the first level DRG to interneuron synapses reduced the number of neurons that would mediate information flow to a single UCoD interneuron (Fig 4G). This interneuron is then connected to 2 interneurons and a single motor neuron.

To again ask how glia could modulate this information flow, we generated a map that labeled tripartite connections and any downstream information flow (Fig 4G). The extent of this could be visualized by color coding the Sankey plot so that synaptic connections with astrocytes and downstream information flow from those are greyed, with information flow without astrocyte control labeled in green (Fig 4G). Although the complexity of the circuit is not reduced as much as synapse thresholding, astroglia control does have the potential to simplify the circuit compared to no synapse thresholding (Fig 4G). To test how astrocyte control could change the interconnectivity, we assumed information flow could be altered by astrocytes. This control would significantly alter the interconnectivity of 1B and 6B DRG neuron’s circuits, shifting the “shared” portion of the circuit by a tripartite synapse located between DRG 1B and an interneuron (Fig S4D). These results revealed an impressive potential for astroglia to modulate information flow within and between two circuits. The potential of glial modulation is similar to increasing the synaptic threshold, highlighting the potential of two cooperative mechanisms to modulate circuit information flow.

The interconnected nature of the connectome predicted that if DRG neurons were simultaneously excited, a large portion of spinal neurons and glia would also be excited or active. To test this, we induced the activity of DRG neurons using cold water immersion and then imaged GCaMP6s as a proxy for neuronal and glial activity^22,44,45^. This sensory assay causes a shivering response by 3 dpf that is dependent on intact afferent axons that connect the DRG neurons with the spinal cord^20,22,45^. This imaging was performed with *Tg(neurod:gal4); Tg(uas:GCaMP6s)* animals at 4 dpf, which express GCaMP6s nearly pan-neuronally in the spinal cord and DRG^44^. We utilized the *neurod* promoter because it still provided sparse labeling that allowed us to identify single neuron cell bodies in the packed spinal cord^44^. As a control to test that the addition of water itself does not cause DRG neuronal activity, we exposed each animal to 23°C water before 4°C water^45^. The dynamics of GCaMP6s does not allow us to detect the order of activity but does indicate if two neurons are likely activated within a similar period of time and within the fluorescent dynamics of GCaMP6s. For each movie, we calculated the z-score of the GCaMP6s intensity over all timepoints and defined an active neuron as a z-score greater than 2.0^44,45^.

The results revealed that in laterally-mounted 5 dpf animals, 81.67 +/-11.67% of individual DRG neurons fired in the first capture immediately following 4^°^C water exposure but were never active after 23^°^C (Fig 4I, S4E). These experiments also revealed that 86.03 +/-4.954% of detectable spinal neurons fired simultaneously with DRG neurons (Fig 4I). Imaging from both lateral and dorsal mounts demonstrated these spinal neurons were responding in both the dorsal and ventral sides of the spinal cord(Fig 4H, Fig S4E), consistent with the locations of interneurons and motor neuron cell bodies. These results were predicted by our serial electron-microscopy mapping of the circuits, which demonstrated a vast network of interneurons that were connected to the DRG neurons in the section (Fig 4A,B, S4A). While most spinal neurons synchronously fired with DRG neurons, 9.527 +/-4.579% (lateral) spinal neurons were not rapidly activated with the DRG neurons(Fig 4I). To provide a readout for contralateral neuronal flow, we also repeated this calcium imaging in dorsally-mounted animals to visualize both sides of the spinal cord (Fig 4H,I). In this analysis, at timepoints in which left-side DRG neurons fired, we detected 71.50 +/-8.249% of spinal neurons rapidly responded to 4^°^C exposure on both the ipsilateral and contralateral side, consistent with our ultrastructural reconstruction (Fig 4H,I).

To test if this spinal neuron activity was propagated from the DRG neurons, we disconnected 8 DRG neurons from the spinal cord circuit via serial axotomies to the centrally-projecting axons of DRG neurons in spinal segments 4-11 (right side) at 5 dpf^22,46,47^, and then repeated the cold water assay at 3 hours post injury (Fig 4J,K). We previously identified that these nerves do not regenerate^22^. When DRG neurons are disconnected, 35.95 +/-16.03% of spinal neurons were rapidly responsive to 4^°^C exposure (Fig 4I,K), demonstrating a significant decrease in the proportion of active spinal neurons, consistent with the idea that spinal neuron activity is dependent on DRG information flow. Despite spinal neuron activity, axotomy did not alter the GCaMP6s activity of DRG neurons (Fig 4K).

Finally, our circuit reconstruction of the glia indicated they had the potential to be integrated into the circuit. To test this possibility, we generated a new transgenic animal, *Tg(gfap:GCaMP6s-CAAX)* that expressed GCaMP6s in astroglia using the regulatory regions of *gfap*^*39*^. We then repeated the cold-water immersion assay and recorded the activity of astroglia in the spinal circuit. GCaMP6s transients were not detected during 23°C exposure but were robustly present after exposure to 4°C water (Fig 4L,M). The robustness of the GCaMP6s signal prevented quantification of single astroglial cells, so we instead quantified the overall volume of GCaMP6s fluorescent changes above threshold as an astroglial population response (Fig 4M). These results demonstrate an immediate increase in GCaMP6s intensity that continued to rise in sequential timepoints (Fig 4M). To determine if this robust response could be predicted by the electron microscopy analysis, we segmented all astroglia located at tripartite synapses in the reconstructed DRG sensorimotor circuits. The GCaMP6s response versus the EM prediction were qualitatively comparable (Fig 4L,N). Together, the functional analysis is consistent with the interconnected nature of individual DRG neuron circuits and the glia within them.

## DISCUSSION

The sensoriomotor circuit is critical for animals to respond to their environment. Here, we mapped the vertebrate sensorimotor circuit at the ultrastructural level. In summary, our work provides a foundational resource of the map of a vertebrate local spinal sensorimotor circuit and introduces a potential layer of modulation that could occur at tripartite synapses. We show that astrocytic processes localize equally to all synapses in simple circuits and that a single astrocyte could regulate multiple and distinct synaptic-connections within a single circuit. This foundational knowledge can be used for future studies that investigate other circuits that underlie important animal behaviors.

Until recent advances in serial electron microscopy and automated cellular reconstructions, circuit maps were typically reserved for model systems with a relatively smaller number of neurons^1^. For example, the *C. elegans* ultrastructural connectome was established in 1986^1^. Even with an organism with 302 neurons, it is clear that the flow of information is complicated. This is particularly the case when animals are exposed to multiple sensory signals, which requires the integration of sensory information. This is exemplified by the neurons in *C. elegans* that respond to harsh vs light touch^48–54^. Both subtypes of neurons synapse onto interneurons in the ventral nerve cord of *C. elegans*, which transmit that information to motor neurons that move muscle^1^. While both subtypes of harsh and light touch neurons are activated during a harsh touch response, the animal responds in a stereotypical way by curling up^51,53–56^. Light touch to the animal however engages a distinct backward or forward movement^56^. Thus, while synapses occur on the same interneuron^1,53,55^, the information of harsh touch is somehow coded to ignore the light touch neuron circuit. How this works in vertebrates is unclear. However, this lack of understanding is not surprising given our incomplete understanding of a vertebrate sensorimotor circuit map.

This level of processing and integration may be expected to be more complex in vertebrates. The sensory neurons in the vertebrate spine are located in the DRG^16,57–59^. The DRG in each somite contains a collection of neurons that each respond to distinct sensory modalities with spatial acuity^57,60–62^. The sensorimotor circuit we mapped contributes to unraveling how this segregation of different sensory inputs can be accomplished. For example, the mapping demonstrated that convergent and divergent circuit connections of individual DRG neurons that could amplify or interrupt information flow. We also observed a network of interconnected interneurons where a single DRG could excite two interneurons, one of which would go on to inhibit an interneuron that was excited by another DRG. The ultrastructural map of these connections is only the first step in understanding how the animal distinguishes an environment that simultaneously activates multiple modalities.

Adding to the complexity of the circuit is the potential role of astrocytes in vertebrates. Studies from invertebrates have already supported the idea that the potential for glia to modulate circuits is evolutionarily conserved^35,36,63^. In *C. elegans*, glial cells control the hyperactivity of dopamine neurons^64^. In Drosophila and zebrafish, astrocytes occupy synapse rich regions and modulate circuits that drive simple behaviors^13,65–68^. We also know that astrocytes in vertebrates can interact extensively with synapses^10^. In mice, it is hypothesized that a given astrocyte can interact with 100K synapses^8,69^. Even more synaptic connections per astrocyte is predicted in humans. We also know that astrocytes provide structural support for neurons and can modulate synaptic function through neurotransmitter regulation^70–72^. In particular, astrocytes express transporters that can remove neurotransmitter from the synapse cleft. They also are hypothesized to participate by releasing gliotransmitters^70,73,74^. However, the extent to which a single circuit can be modulated by astrocytes needs more investigation. Our work shows the limits of the potential for astrocytic modulation at the synaptic level in a vertebrate sensorimotor circuit. In our map, we revealed that astrocytic processes contact synapses equally throughout the circuit. There are several reasons why such distribution would be advantageous. It is possible that synaptic control at each area of the circuit ensures that there are multiple levels that can be controlled, including initial relay of sensory information, integration of information from interneurons and finally firing of motor neurons to drive muscle movement. The equal distribution of tripartite synapses also might reflect the development of astrocytes and synapses, which is still occurring at 6 dpf when the EM dataset was collected. These concepts are beyond the scope of this report but warrant future exploration.

The connectome of the nervous system is a critical and foundational resource if we want to further investigate the development and functionality of circuits. This work provides a key exploration of the connectivity of the vertebrate sensorimotor circuit and reveals the precise synaptic locations where glial cells could modulate the circuit.

## LIMITATIONS

The circuit map includes only chemical synapses and thereby neglects the important contribution of electrical synapses in information flow. The circuit map is generated from a 6 dpf zebrafish and thus likely represents a circuit that will change as the animal ages. Further, the full complex connectivity of local spinal sensorimotor circuits is likely to expand significantly beyond what is reported here because synaptic connections outside of the 3 hemisegments are not included. Similarly, the represented circuit is likely to expand in animals with a larger number of complex neurons. The number of tripartite synapses is likely underestimated because of current approaches that limit the segmentation of small astrocytic processes. Further, this report does not consider the potential astrocytic modulation of information flow beyond direct contact with synaptic cleft regions, which is well reported in the literature but can be the focus of future investigation^13,75,76^.

## METHODS

### Experimental Model and subject details

Experiments procedures followed the NIH guide for the care and use of laboratory animals. The Animal studies were approved by University of Notre Dame Institutional Animal Care and Use Committee (IACUC) (protocol 19-08-5464), which is guided by the United States Department of Agriculture, the Animal Welfare Act (USA) and the Assessment and Accreditation of Laboratory Animal Care International.

Animal Specimens. *Danio rerio* (zebrafish) were utilized in this study. The stable strains used for this study were: AB, *Tg(UAS:GCaMP6s)*^*77*^, *Tg(neurod:Gal4+myl7:GFP)*^*78*^, *Tg(gfap:GCaMP6s-CAAX)*(generated for this study), *Tg(sox10:mRFP)*^*79*^. All embryos were produced through pairwise matings and grown in 28°C in constant darkness. At 24 hpf, zebrafish were exposed to PTU (0.0003%) to reduce pigmentation for intravital imaging. Age of animals was determined by hour post fertilization and stages of development^80^.

### Experimental Procedures

#### Generation of transgenics

*Tg(gfap:GcaMP6s-CAAX)* was created by injecting tol2 RNA and psSL08 (*gfap*:GcaMP6s-CAAX-pA) into one cell animals. The plasmid was generated using gateway plasmids (p5e-*gfap*, pMe-GcaMP6s-CAAX, p3e-polyA, p394). Injected animals were grown to adults and then screened for GCAMP6s-CAAX fluorescence. Once founders were identified, animals were outcrossed to generate a stable *Tg(gfap:GcaMP6s-CAAX)* transgenic.

#### Serial electron microscopy

The serial electron microscopy dataset used in this study was previously published in Svara et al. 2018^23^. The images were collected with serial block-face electron microscopy from a 6 day post fertilization zebrafish larva. The images were collected from the middle of the spinal cord, near the anal pore. The volume is 74 × 74 × 207 µm^3^, with a voxel size of 9 × 9 × 21 nm^3^. All the EM images for this study were managed using https://knossos.app^24^.

#### Semi-automated cellular reconstructions

Cellular reconstructions in this study were completed using Knossos based on Svara et al. 2022^24^. Semiautomated reconstructions were completed by clicking on the cell of interest in Knossos. This revealed the morphology of the cell in the reconstruction window. To complete the cellular reconstructions, each endpoint was confirmed. If the axon continued in the section, segments of neurons were joined using the agglomeration function in Knossos. Cells were reconstructed until all endpoints were confirmed. For a subset of reconstructions, improper mergers of the agglomeration prevented the semiautomated approach. These neurons were reconstructed using the skeletonization function in Knossos. The center of the axon was marked throughout the section. The skeletonization reconstruction included nodes every 10 z-positions in the section. All skeletonized neurons were confirmed with two authors. Glial cells were reconstructed with the semi-automated approach.

#### Automated Synaptic Identification

Synapses were identified using the procedures outlined in Svara et al. 2022 and reconstructed using the 3dEMtrace service provided by ariadne.ai ag (https://ariadne.ai)^24^. Synapses were defined as: 1. Two adjacent cellular surfaces were parallel and juxtaposed. 2: Vesicle clouds were present in the axon in close proximity to the membrane and opposite the adjacent cell. 3: A thick and dark labeling, although faint at times, of the postsynaptic membrane was present^24^. Two parts of the synapse were labeled throughout the dataset. Characteristics 1 and 3 were marked by pink and synaptic clouds defined by characteristic 2 were labeled in blue. We defined mature synapses as requiring both pink and blue markings and thereby all three characteristics. While synapses without all their characteristics were initially mapped for sensory neurons, the remainder of the mapping only included synapses with all three characteristics.

#### Neuron classification

Neurons were classified based on their morphology and previous literature that defined different subtypes of cells in the spinal cord. A decision tree was created based on the previous literature and each cell was classified by the newly created decision tree. The excitatory vs inhibitory classification was based on cross-referencing the neuron morphologies with neurotransmitter classification in published literature ^19,25–29^. CoSA neurons were excluded from analysis with excitatory/inhibitory because they are characterized in the literature as both.

#### Glia classification

Astroglia subtypes were defined by the morphology of the cell. Cells with radial projections that extended to the edge of the spinal cord and made up a significant portion of the glial limitans were characterized as glial limitans astroglia. Protoplasmic astroglia exhibited morphology with numerous small processes that extended into the neuropil. Glia that exhibit large mergers were not included in the subclassification. This represented 24.42% of astroglia that were present at synapses. For visualization in Fig 3E, improper mergers between astroglia and neurons were removed via Photoshop. For transparency’s sake, unedited and edited versions are represented in Fig S3.

#### Tripartite Synapses

At every defined synapse (per the criteria above), all membranes directly in contact with both pre and post synaptic membranes were clicked to display the cell reconstruction and identify it as astroglia. Any “tertiary” membrane that revealed an astrocyte through semi-automated reconstructions was characterized as a tripartite synapse.

#### *In vivo* imaging

Animals were anesthetized with veterinary grade 3-aminobenzoic acid ester during mounting and then were bathed in PTU during imaging^21,22,44,46,78^. A glass-bottom 35 mm petri dish was used to image the animals. Each animal was mounted on the glass-bottom dish with 0.8% low melt agarose. All confocal images were collected on a custom-built (3i) spinning disk confocal microscope. The microscope contains: Zeiss Axio Observer Z1 Advanced Mariana Microscope, X-cite 120LED White Light LED System, filter cubes for GFP and mRFP, a motorized X,Y stage, piezo Z stage, 20X Air (0.50 NA), 63X (1.15NA), 40X (1.1NA) objectives, CSU-W1 T2 Spinning Disk Confocal Head (50 µm) with 1X camera adapter, and an iXon3 1Kx1K EMCCD camera or Prime 95B back illuminated CMOS camera, dichroic mirrors for 446, 515, 561, 405, 488, 561,640 excitation, laser stack with 405 nm, 445 nm, 488 nm, 561 nm and 637 nm. Z-stacks were collected with a step size of 1 um covering the spinal cord of the animal. Brightness and contrast were only adjusted for generation of the figures in this study.

#### Temperature Submersion and GCaMP6s assays

Animals were exposed to PTU from 1-5 dpf to restrict skin pigmentation. All imaging was performed on the spinning disk confocal inverted microscope as noted in microscopy section. *Tg(gfap:GCaMP6s-CAAX)* larvae were raised until 5 dpf for analysis of astroglia. The *Tg(gfap:GCaMP6s-CAAX)* animals were mounted laterally, and the glial limitans/spinal cord boundary was positioned 2-4 um from the bottom of a 40 um z-stack (2 um step-size) in order to fully capture the glia of the spinal cord.

*Tg(neurod:gal4+myl7-GFP);(UAS:GCaMP6s)* animals were raised and imaged at 5 dpf to image neuronal activity. For the neuronal activity experiments, both laterally and dorsally mounted *Tg(neurod:Gal4+myl7-GFP);(uas:GCaMP6s)+* animals were used. In either mounting position, the appearance of the DRG was positioned 5-7 microns from the bottom of a 40 um z-stack (2 micron step-size).

In the injury context, *Tg(deurod:gal4+myl7-GFP);(uas:GCaMP6s);Tg(sox10:mRFP)* animals were raised to 5 dpf at which point the centrally-projecting axons at spinal segment 4-11 were injured as previously described^22,46^; 3 hours post injury, these animals underwent the cold water immersion assay during calcium imaging^20,22,44^.

For the water immersion assay, animals were anesthetized and individually mounted in 0.8% low melting agarose in 4-well glass bottom dishes, wherein the agar was thin enough that the body of the animal created a bump in the agar. After the agar solidified, fresh, room temperature egg water was added to the wells and the animals were given 20 minutes to recover from the anesthetic. Egg water was then removed from the well and a 24-time point timelapse was started. At a step size of 2 um, z-range of 40 um, and exposure time set to 85 ms, each capture of the timelapse took exactly 5-sec before the next capture began. During time point 7, 23°C egg water was added to the well, just before the 8th time point. Beginning after time point 11, the water is carefully aspirated from the well without disturbing the dish/the xy position. At time point 15 - just before timepoint 16, 4°C egg water was added to the well, where it remained for the rest of the 24 timepoint timelapse movie.

#### Sankey Plots

Cells were given individual cell identity numbers in Knossos based on the segmentation ID of the neuronal cell body. A custom code was written to generate sankey plot information. In brief, each the segmentation IDs of each synaptic connection in the circuit was generated. For example, segmentationID1 [2] segmentationID2 would represent cell#1 is connected through 2 synapses to cell#2. This process was completed for every connection in the circuit. Once completed for a single DRG neuron’s circuit, the code was repeated on each circuit. To represent the interconnectivity of the circuits, synaptic connections between DRG 1B and 6B were combined. Sankey plot data was then imported into sankeymatic and color coded according to the figure panel.

#### Quantifications and Statistical Tests: Quantification of cell measurements

The area of the axon was measured using the line ROI tool in ImageJ. The total area of the ROI was first calculated in pixels and then converted to um. The swelling at the synapse was measured where the synaptic cloud are centered. Before and after the swellings were defined as directly adjacent locations from the synaptic cloud regions. Length of DRG axons was estimated by taking multiple points within the axons and summing the Euclidean distance between each of these points, with respect to the scalebar given in Knossos.

#### Quantification of synaptic and glia organization

Synapses and glia at synapses were counted within each DRG neuron’s circuit. To keep the data consistent across DRG circuits, all the data was combined into a larger database. 3 lists were made using a custom Python code – an object list, a synapse list and a tripartite glia list. Each element of the object list had 2 components – the Knossos ID of the object, and the cell type of the object. Each element of the synapse list had 4 components – the coordinate of the synapse, the synaptic type, the pre-synaptic reference number, and the post-synaptic reference number (pointing to the position of the objects within the object list). Each element of the tripartite glia list had 3 components – the coordinate of the glia, the Knossos ID of the glia, and the synaptic reference number (pointing to the position of the synapse within the synapse list). These lists were populated with data from manually counting the synapses, first the object list, then the synapse and tripartite glia list together. Duplicate synapses across circuits were removed by getting the Euclidean distance between known coordinates and setting a threshold of 35 voxels, which we determined a distance that ensured unique synapses could be identified.

#### Quantification of synapse types

Synapse types were identified into 4 subtypes by where the synapse occurred with respect to the pre-synaptic and post-synaptic neurons: A-A for axon to axon, A-D for axon to dendrite, A-S for axon to soma, and the rare A-G for axon to glia. Axons, dendrites and cell somas were defined by specific criteria that define that area of the neuron. Cell bodies were defined by the location of the nucleus. Neuronal branches were defined as dendrites if they extended multiple projections from the cell soma and branched extensively into the neuropil. Dendrites typically did not contain synaptic clouds/vesicles. Axons were defined as singular processes that extended from the cell soma and contained pre-synaptic material.

#### Quantification of tripartite types

Glial modulators were identified by morphology as mentioned above (under Tripartite Synapses), and the number of unique glia and the number of times each one appeared in a given circuit were calculated.

#### Quantification of GCaMP6s co-activity

Maximum z-projections were made of each capture. GCaMP6s was analyzed by exporting the 16-bit timelapse movies into ImageJ software (Fiji). A single rectangle was drawn around any apparent fluorescent/cellular landmarks that could be distinguished throughout the timelapse. Using the Template Matching plugin, frames of each timelapse were aligned according to the visible landmarks throughout the timelapse to correct for any movements from the animal and/or additions of water.

For the glial calcium imaging, the scale of the file is set according to the microscope’s specifications, in which the distance in pixels is scaled to make a 1.0 pixel aspect ratio and known distance of 1.00 microns. The threshold plugin was run and Otsu settings were used to make the imaging file binary. The low-end threshold is set so that almost no ROIs are visible in timepoints 1-7, when no calcium activity would be expected. The high end of the threshold was set at its highest setting. The Analyze Particles feature was then used with the size was set to 3.00-Infinity (microns^ 2). It was then confirmed that no spots were identified during frames 1-7 unless a random activation event occurred. This process was completed for each timelapse, however, the low-end threshold values varied in each animal. From each of the 24 timepoints, the average and standard deviation was used to calculate the z-score of the integrated density (the product of area and mean fluorescent value) in each frame. A z-score of 2.0 or more was set as a significant activation event. These z-scores were plotted into a stacked xy line graph or heat map. The total area above threshold was also used to display the global activation of the glial limitans.

For the neuronal calcium imaging, the first capture immediately after the cold water immersion was used to set ROIs. ImageJ’s ROI Manager feature was utilized and all distinguishable (regardless of visible brightness) GCaMP+ DRG and spinal neurons were freehand traced and added to the ROI manager. Once all cells were traced in 4°C immersion timepoint, any other neurons not previously traced that fired in later time points were also traced. The multi-measure tool was used to obtain the integrated density for each traced cell. The integrated density measurements for each cell at each timepoint was used to obtain the average and standard deviation in order to calculate z-scores. A z-score of 2.0 or more was marked as a neuronal firing event. Neuronal GCaMP6s dynamics were classified into 3 categories. If GCaMP6s changes underwent a z-score change of >2.0 in the 16th frame it was classified as a rapid firing event. If a neuron did not have a z-score of 2.0 or more until the 17th frame or later, that was classified as a delayed firing event. If the cell never underwent a z-score change of 2.0 or more then that was classified as no firing event. z-scores for all neurons were plotted in a stacked xy line graph for each animal. The percent of rapid, delayed and no firing event spinal neurons was collected for each animal for comparisons. A Fisher’s exact test was used to compare the percentage of rapidly responding versus delayed firing spinal neurons between groups. Note, in the injured GCaMP+ animals, only DRG 6-9 were analyzed to ensure that all spinal neurons analyzed in this assay were flanked by at least two DRG that had previously been injured.

#### Statistical analysis

Prism was used for all statistical analysis. Sample sizes were based on previous publications but were not predetermined by statistical methods. All statistical tests were performed with biological replicates and not technical replicates. Some analysis excluded any data points associated with neuronal branches without cell soma in the EM section. Heatmaps of astrocyte synaptic interactions excluded astrocytes that were merged. Data extracted from EM were from a single healthy animal. For GCaMP6s experiments, healthy animals were randomly selected. GCaMP6s experiments were repeated at least twice.

#### Software

Slidebook, Prism, ImageJ, Adobe Illustrator, and Knossos were used to acquire, analyze, and compile figures. Sankey plots were generated with https://sankeymatic.com/build/.

#### Custom coding

Custom code was written for Python and distributed in github (https://github.com/ZacharyMKoh/Spinal-Sensorimotor-Circuit). Custom code was used to generate Figures 1E, 1H, 2C, 2E, 2F, 2G, 2H, 2I, 3B, 3C, 3G, 4A, 4B, 4C, 4D, 4E, 4F, 4G, S1B-D, S1I-L, S2C, S3A, S4A, S4C-D. Data generated from the custom code was imported into Prism or sankeymatic for figure generation.

## DATA AVAILABILITY

All data collected for this study are included in the figures and supplementary material.

## ACKNOWLEDGEMENTS

We thank members of the Smith lab for helpful discussions. Thank you to ariadne for field questions regarding cellular reconstructions, Johann Bollmann and Winfried Denk for their contributions to the serial EM dataset, and David Lyons for sharing *Tg(UAS:GCaMP6s)* transgenic zebrafish. We also thank 3i for fielding imaging questions and Deborah Bang and others for maintaining the zebrafish facility. This work was supported by The University of Notre Dame College of Science Summer Research Fellowship (KAA, KC, ZK), Michael and Elizabeth Gallagher Family (CJS), The University of Notre Dame (CJS), the SMART foundation (CJS), and the NIH (DP2NS117177)(CJS). The funders had no role in study design, data collection, analysis, decision to publish or preparation of the manuscript.

## AUTHOR CONTRIBUTIONS STATEMENT

Ricky Avalos Arceo, Khang Chau, Zachary M. Koh, Antonio Dolojan, Jacob Hammer, and Sarah E.W. Light performed the analysis and experimentation and wrote and edited the manuscript.

Ricky Avalos Arceo supervised the reconstruction efforts.

Zachary M. Koh generated all code for analysis

Fabian Svara, Michał Januszewski, provided the serial electron microscopy dataset and the segmentation.

Fabian Svara (ariadne.ai ag) provided the automated synaptic identification

Cody J. Smith performed analysis, conceived, funded, and supervised the project, and wrote and edited the manuscript.

## COMPETING INTERESTS STATEMENT

Fabian Svara is employed by and holds shares in ariadne.ai ag. Michał Januszewski is employed by Google. The remaining authors declare no competing interests.

## SUPPLEMENTAL MATERIAL

**Figure S1.**
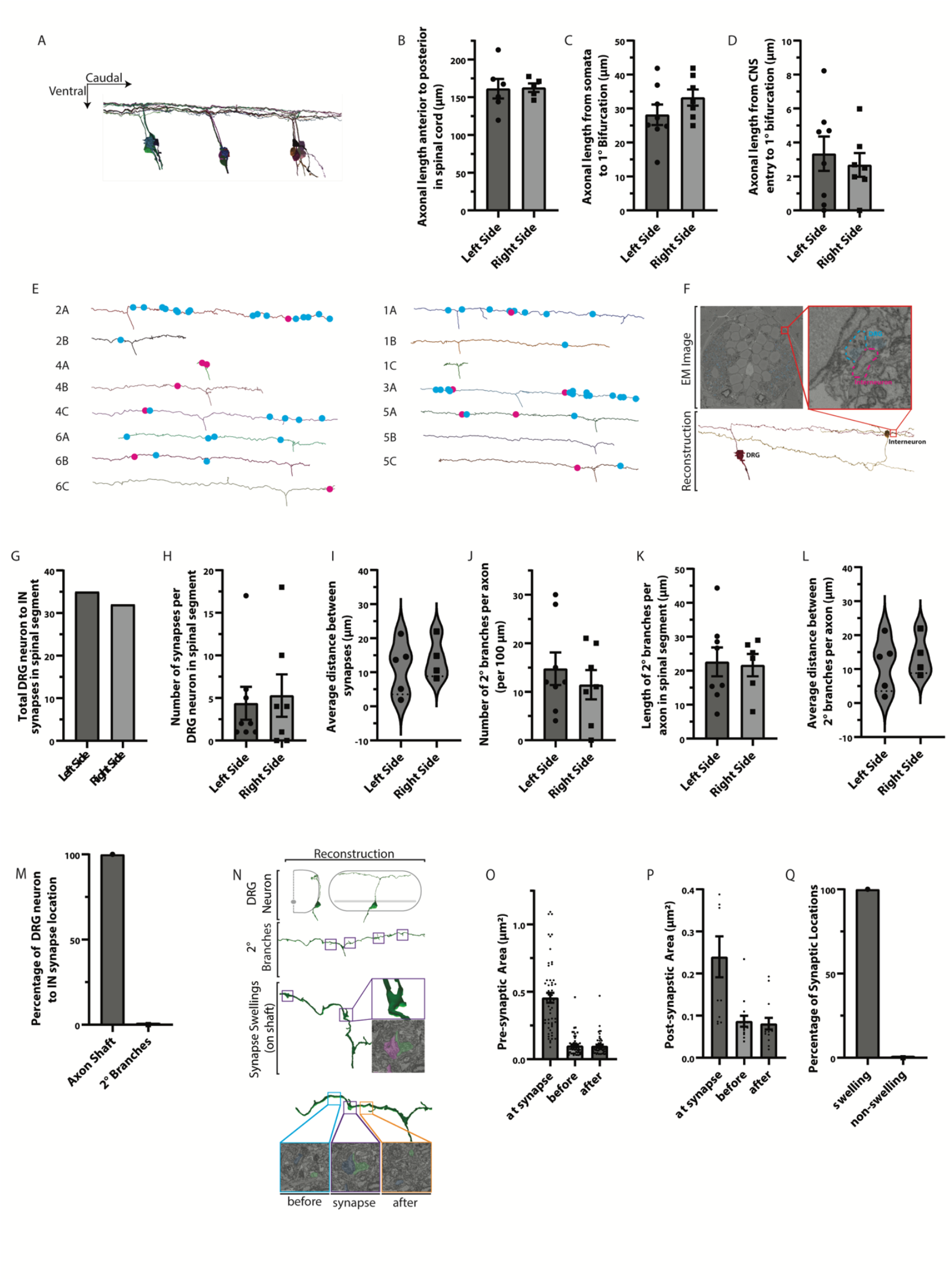
Reconstruction of sensorimotor circuit. A. Segmentation reconstruction of the 6 dpf zebrafish DRG neurons in a lateral view. B-D. Quantification of DRG axonal length in the spinal cord (B), from soma to axon bifurcation (C) and from the CNS entry site to bifurcation (D). E. Reconstruction of individual DRG neurons and their synaptic sites. Blue dots denote pre-synaptic sites, pink dots denote post-synaptic sites. F. EM image and reconstructions of a synaptic connection between a DRG neuron and interneuron. Inset shows synapse with pre-synaptic DRG outlined in blue and post-synaptic interneuron outlined in pink. G. Quantification of the total number of DRG to interneuron (IN) synapses in the tissue section. H-M. Quantification of the number of synapses per DRG neuron in the spinal segment (H), distance between synapse sites (I), number of secondary branches on DRG neurons (J), length of secondary DRG branches (K), average distance between secondary branches (L), and percentage of DRG neuron to interneuron synapses that occur at secondary branches vs the axon shaft (M). N. Reconstruction of a DRG neuron showing swellings along that axon where synaptic sites are located. O-Q. Quantification of the axonal shaft area of DRG neurons at synaptic swellings, and before and after swellings (O), the dendritic shaft area at synapse, before, and after swellings (P), and the percentage of synaptic locations at swelling vs non-swellings (Q). T-test: B-D, H-L, O-P. Fishers Exact test: M, Q.

**Figure S2.**
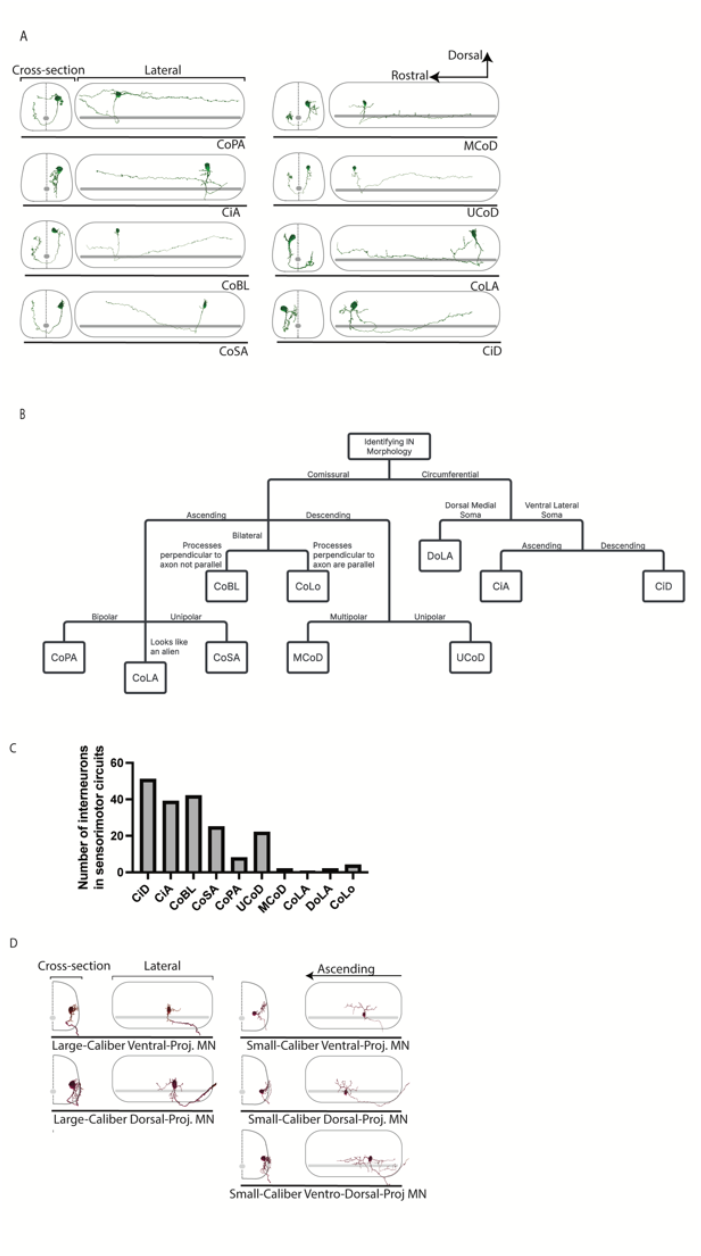
Classes of neuron subtypes within the DRG circuits. A. Cellular reconstructions of neuron subtypes based on morphology in the EM section. Dorsal up and rostral left. Traverse and lateral cross-sections are shown. B. Decision tree to determine the interneuron class. C. Quantification showing the number of interneurons of each subtype in the local spinal sensorimotor circuit. D. Cellular reconstructions of motor neurons that are connected in the sensorimotor circuit showing different morphotypes of motor neurons.

**Figure S3.**
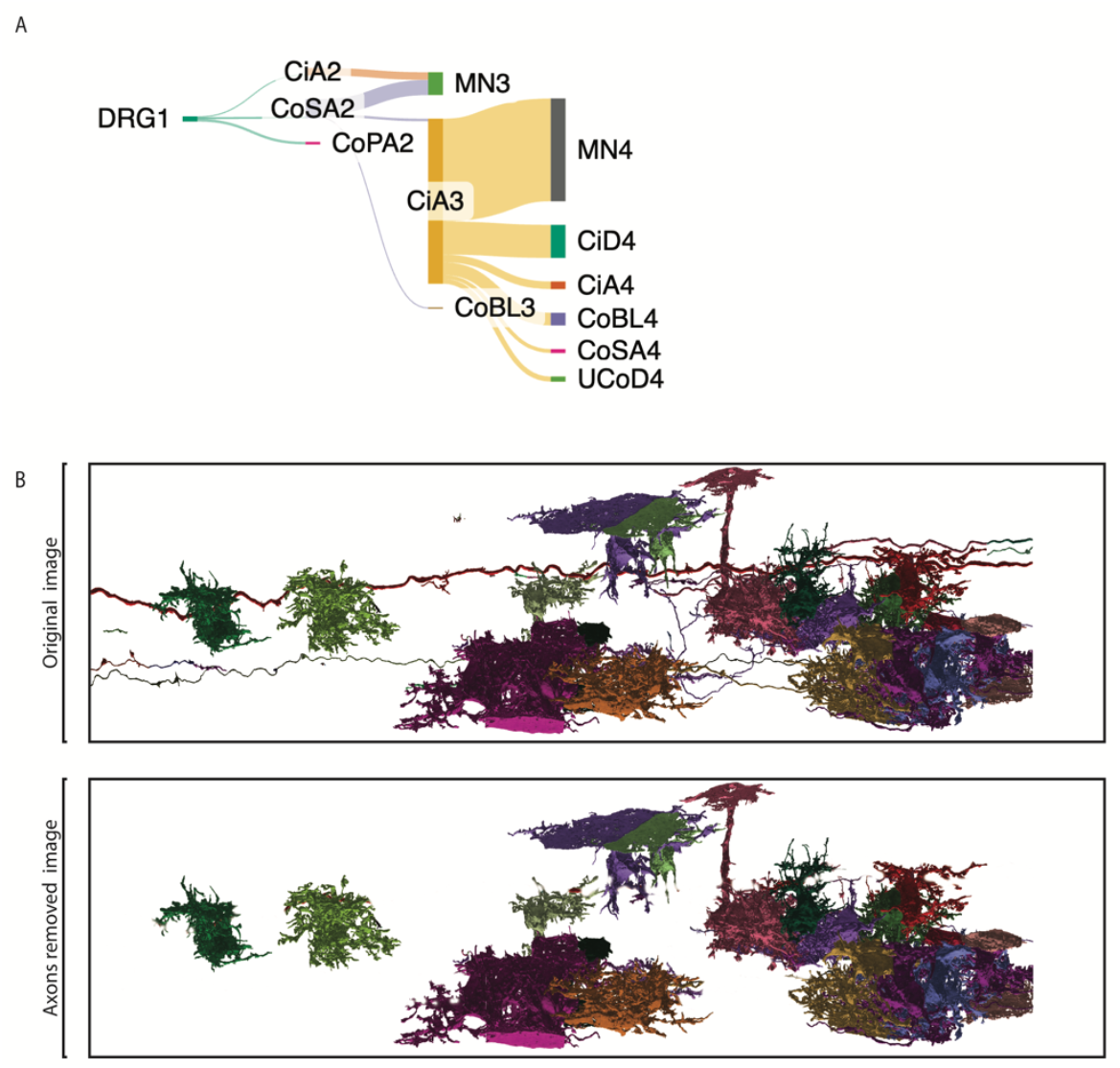
Glia associated with a single DRG neuron’s circuit. A. Schematic showing potential information flow demonstrating each of the classes of neurons in the circuit represented in Fig 3. Circuit is the same as shown in Fig 3C. B. Cellular reconstructions of astroglia that associate with a single DRG neuron’s circuit in Figure 3. The original image is the raw reconstructions that are downloaded from the reconstruction server showing that some astroglia are improperly merged with neurons. The improper axons were removed from the image to provide a more accurate reconstruction of the astrocytes. For transparency, both images are presented.

**Figure S4.**
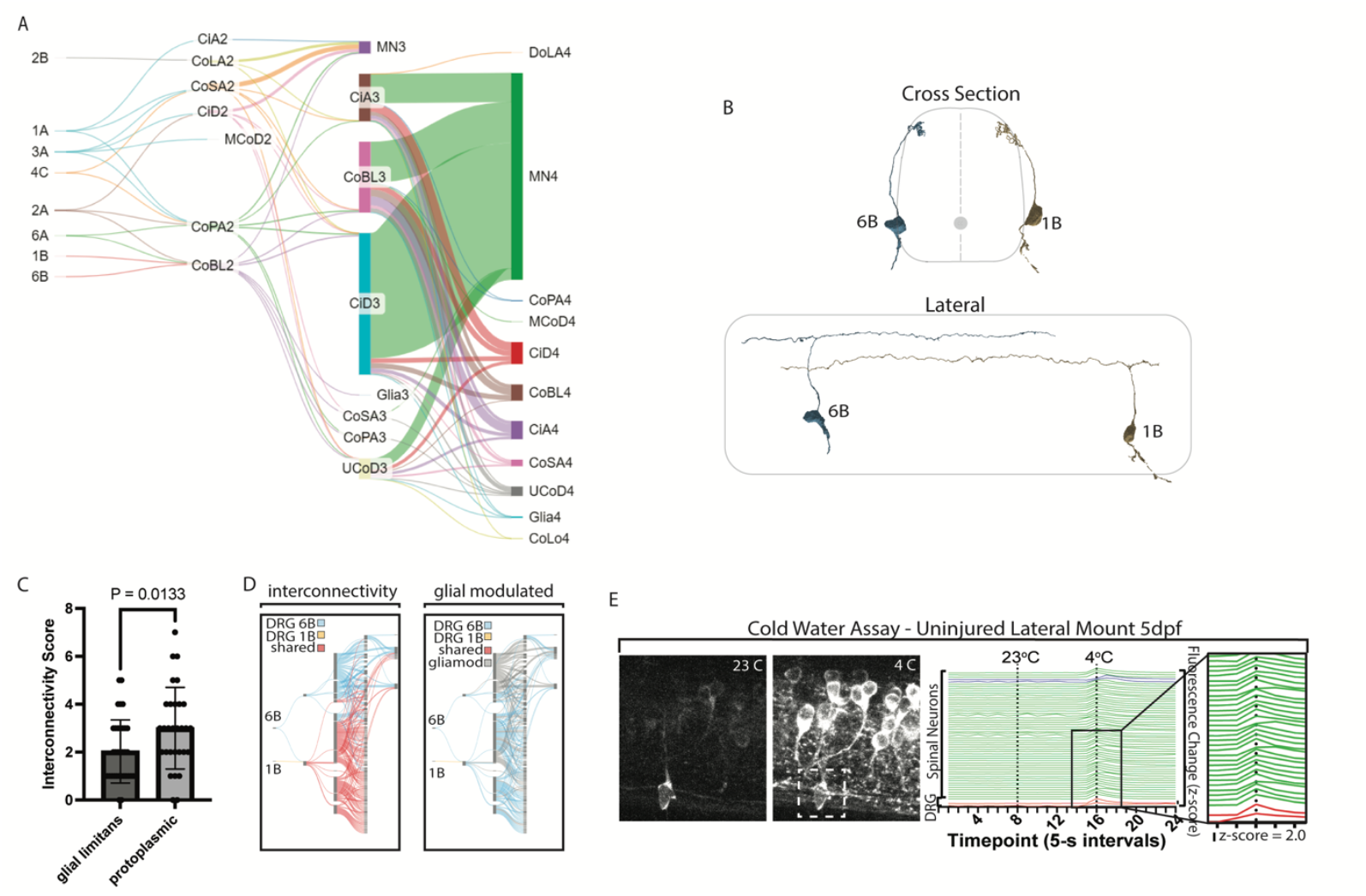
Potential flow of information through DRG circuits. A. Sankey plot showing the flow of information from DRG neurons to interneurons and then to motor neurons. The width of the bar represents the number of synapses for that given class of neuron. Colors are randomly assigned. B. Reconstructions of DRG 1B and 6A that are used as examples of interconnected circuits in Fig 4. Note that the DRG are on opposite sides of the spinal cord. C. Quantification of the interconnectivity score, which is defined as the number of circuits with which a given astrocyte is connected. Note the protoplasmic astrocytes are more connected than glial limitans astroglia. D. Sankey plot showing the interconnectivity of DRG 1B and 6A and how that changes if glia could modulate the circuit. Synaptic connections with glia contacts and their downstream connections are gray. E. Confocal images and quantification of DRG and spinal neurons when *Tg(neurod1:Gal4); Tg(UAS:GCaMP6s* animals are exposed to 23°C and 4°C water and mounted laterally. z-score changes of intensity are shown. DRG neurons with a rapid change in z-score >2.0 are colored red, spinal neurons with rapid changes are colored green, spinal neurons with delayed changes are blue, and neurons that do not change (z-score>2.0) are black. Exposure to 4°C water causes a rapid increase in DRG and spinal neuron GCaMP6s intensity.

## REFERENCES

1. JG, W., E, S., JN, T., and S, B. (1986). The structure of the nervous system of the nematode Caenorhabditis elegans. Philos Trans R Soc Lond B Biol Sci 314, 1–340. 10.1098/RSTB.1986.0056.

2. Abbott, L.F., Bock, D.D., Callaway, E.M., Denk, W., Dulac, C., Fairhall, A.L., Fiete, I., Harris, K.M., Helmstaedter, M., Jain, V., et al. (2020). The Mind of a Mouse. Cell 182, 1372–1376. 10.1016/j.cell.2020.08.010.

3. Pospisil, D.A., Aragon, M.J., Dorkenwald, S., Matsliah, A., Sterling, A.R., Schlegel, P., Yu, S., McKellar, C.E., Costa, M., Eichler, K., et al. (2024). The fly connectome reveals a path to the effectome. Nature 634, 201–209. 10.1038/s41586-024-07982-0.

4. Winding, M., Pedigo, B.D., Barnes, C.L., Patsolic, H.G., Park, Y., Kazimiers, T., Fushiki, A., Andrade, V., Khandelwal, A., Valdes-Aleman, J., et al. (2023). The connectome of an insect brain. Science (1979) 379. 10.1126/science.add9330.

5. Schlegel, P., Yin, Y., Bates, A.S., Dorkenwald, S., Eichler, K., Brooks, P., Han, D.S., Gkantia, M., dos Santos, M., Munnelly, E.J., et al. (2024). Wholebrain annotation and multi-connectome cell typing of Drosophila. Nature 634, 139–152. 10.1038/s41586-024-07686-5.

6. Dorkenwald, S., Matsliah, A., Sterling, A.R., Schlegel, P., Yu, S., McKellar, C.E., Lin, A., Costa, M., Eichler, K., Yin, Y., et al. (2024). Neuronal wiring diagram of an adult brain. Nature 634, 124–138. 10.1038/s41586-024-07558-y.

7. Lin, A., Yang, R., Dorkenwald, S., Matsliah, A., Sterling, A.R., Schlegel, P., Yu, S., McKellar, C.E., Costa, M., Eichler, K., et al. (2024). Network statistics of the whole-brain connectome of Drosophila. Nature 634, 153–165. 10.1038/s41586-024-07968-y.

8. Halassa, M.M., Fellin, T., Takano, H., Dong, J.H., and Haydon, P.G. (2007). Synaptic islands defined by the territory of a single astrocyte. Journal of Neuroscience 27, 6473–6477. 10.1523/JNEUROSCI.1419-07.2007.

9. Clarke, L.E., and Barres, B.A. (2013). Emerging roles of astrocytes in neural circuit development. Preprint at Nat Rev Neurosci, 10.1038/nrn3484 10.1038/nrn3484.

10. Chung, W.S., Allen, N.J., and Eroglu, C. (2015). Astrocytes Control Synapse Formation, Function, and Elimination. Cold Spring Harb Perspect Biol 7. 10.1101/CSHPERSPECT.A020370.

11. Stobart, J.L., Ferrari, K.D., Barrett, M.J.P., Gl, C., Stobart, M.J., Zuend, M., Weber, B., Stobart, M.J., and Zuend, M. (2018). Cortical Circuit Activity Evokes Rapid Astrocyte Calcium Signals on a Similar Timescale to Neurons. Neuron, 1–10. 10.1016/j.neuron.2018.03.050.

12. Takano, T., Wallace, J.T., Baldwin, K.T., Purkey, A.M., Uezu, A., Courtland, J.L., Soderblom, E.J., Shimogori, T., Maness, P.F., Eroglu, C., et al. (2020). Chemico-genetic discovery of astrocytic control of inhibition in vivo. Nature. 10.1038/s41586-020-2926-0.

13. Mu, Y., Bennett, D. V., Rubinov, M., Narayan, S., Yang, C.-T., Tanimoto, M., Mensh, B.D., Looger, L.L., and Ahrens, M.B. (2019). Glia Accumulate Evidence that Actions Are Futile and Suppress Unsuccessful Behavior. Cell, 1–17. 10.1016/j.cell.2019.05.050.

14. Ladle, D.R., Pecho-Vrieseling, E., and Arber, S. (2007). Assembly of Motor Circuits in the Spinal Cord: Driven to Function by Genetic and Experience-Dependent Mechanisms. Neuron 56, 270–283. 10.1016/j.neuron.2007.09.026.

15. Meltzer, S., Santiago, C., Sharma, N., and Ginty, D.D. (2021). The cellular and molecular basis of somatosensory neuron development. Neuron 109, 3736–3757. 10.1016/j.neuron.2021.09.004.

16. Cajal, S.R. (1911). Histology of the nervous system of man and vertebrates (1909{\textendash}1911), vols. 1 and 2 (L. Azoulay, Spanish trans.; N. Swanson and LW Swanson, French∼…. Preprint.

17. Harding, E.K., Fung, S.W., and Bonin, R.P. (2020). Insights Into Spinal Dorsal Horn Circuit Function and Dysfunction Using Optical Approaches. Front Neural Circuits 14. 10.3389/fncir.2020.00031.

18. McGraw, H.F., Nechiporuk, A., and Raible, D.W. (2008). Zebrafish Dorsal Root Ganglia Neural Precursor Cells Adopt a Glial Fate in the Absence of Neurogenin1. Journal of Neuroscience 28, 12558–12569. 10.1523/JNEUROSCI.2079-08.2008.

19. Lewis, K.E., and Eisen, J.S. (2003). From cells to circuits: development of the zebrafish spinal cord. Prog Neurobiol 69, 419–449. 10.1016/S0301-0082(03)00052-2.

20. Kikel-Coury, N.L., Green, L.A., Nichols, E.L., Zellmer, A.M., Pai, S., Hedlund, S.A., Marsden, K.C., and Smith, C.J. (2021). Pioneer Axons Utilize a Dcc Signaling-Mediated Invasion Brake to Precisely Complete Their Pathfinding Odyssey. The Journal of Neuroscience 41, 6617–6636. 10.1523/JNEUROSCI.0212-21.2021.

21. Nichols, E.L., and Smith, C.J. (2019). Pioneer axons employ Cajal’s battering ram to enter the spinal cord. Nat Commun 10. 10.1038/s41467-019-08421-9.

22. Nichols, E.L., and Smith, C.J. (2020). Functional Regeneration of the Sensory Root via Axonal Invasion. Cell Rep 30, 9-17.e3. 10.1016/j.celrep.2019.12.008.

23. Svara, F.N., Kornfeld, J., Denk, W., and Bollmann, J.H. (2018). Volume EM Reconstruction of Spinal Cord Reveals Wiring Specificity in Speed-Related Motor Circuits. Cell Rep 23, 2942–2954. 10.1016/j.celrep.2018.05.023.

24. Svara, F., Förster, D., Kubo, F., Januszewski, M., Maschio, M., Schubert, P.J., Kornfeld, J., Wanner, A.A., Laurell, E., Denk, W., et al. (2022). Automated synapse-level reconstruction of neural circuits in the larval zebrafish brain. 19. 10.1038/s41592-022-01621-0.

25. Satou, C., Kimura, Y., Kohashi, T., Horikawa, K., Takeda, H., Oda, Y., and Higashijima, S. (2009). Functional role of a specialized class of spinal commissural inhibitory neurons during fast escapes in zebrafish. J Neurosci 29, 6780–6793. 10.1523/JNEUROSCI.0801-09.2009.

26. Batista, M.F., Jacobstein, J., and Lewis, K.E. (2008). Zebrafish V2 cells develop into excitatory CiD and Notch signalling dependent inhibitory VeLD interneurons. Dev Biol 322, 263–275. 10.1016/j.ydbio.2008.07.015.

27. Higashijima, S., Schaefer, M., and Fetcho, J.R. (2004). Neurotransmitter properties of spinal interneurons in embryonic and larval zebrafish. Journal of Comparative Neurology 480, 19–37. 10.1002/cne.20279.

28. Wilson, A.C., and Sweeney, L.B. (2023). Spinal cords: Symphonies of interneurons across species. Front Neural Circuits 17. 10.3389/fncir.2023.1146449.

29. Björnfors, E.R., and El Manira, A. (2016). Functional diversity of excitatory commissural interneurons in adult zebrafish. Elife 5. 10.7554/eLife.18579.

30. Babin, P.J., Goizet, C., and Raldúa, D. (2014). Zebrafish models of human motor neuron diseases: Advantages and limitations. Prog Neurobiol 118, 36–58. 10.1016/j.pneurobio.2014.03.001.

31. D’Elia, K.P., Hameedy, H., Goldblatt, D., Frazel, P., Kriese, M., Zhu, Y., Hamling, K.R., Kawakami, K., Liddelow, S.A., Schoppik, D., et al. (2023). Determinants of motor neuron functional subtypes important for locomotor speed. Cell Rep 42, 113049. 10.1016/j.celrep.2023.113049.

32. Bereciartu, H. Pío del Río-Hortega: The Revolution of Glia. 10.1002/ar.24266.

33. Chen, J., Poskanzer, K.E., Freeman, M.R., and Monk, K.R. (2020). Live-imaging of astrocyte morphogenesis and function in zebrafish neural circuits. Nat Neurosci 23, 1297–1306. 10.1038/s41593-020-0703-x.

34. Chen, J., Stork, T., Kang, Y., Nardone, K.A.M., Auer, F., Farrell, R.J., Jay, T.R., Heo, D., Sheehan, A., Paton, C., et al. (2023). Astrocyte growth is driven by the Tre1/S1pr1 phospholipid-binding G protein-coupled receptor. Neuron 0. 10.1016/J.NEURON.2023.11.008.

35. Freeman, M.R. (2015). Drosophila central nervous system glia. Cold Spring Harb Perspect Biol 7. 10.1101/cshperspect.a020552.

36. Freeman, M.R. (2010). Specification and morphogenesis of astrocytes. Science (1979) 330, 774–778. 10.1126/science.1190928.

37. Freeman, M.R., Delrow, J., Kim, J., Johnson, E., and Doe, C.Q. (2003). Unwrapping glial biology: Gcm target genes regulating glial development, diversification, and function. Neuron 38, 567–580. 10.1016/S0896-6273(03)00289-7.

38. Johnson, K., Barragan, J., Bashiruddin, S., Smith, C.J., Tyrrell, C., Parsons, M.J., Doris, R., Kucenas, S., Downes, G.B., Velez, C.M., et al. (2016). Gfap-positive radial glial cells are an essential progenitor population for later-born neurons and glia in the zebrafish spinal cord. Glia 64, 1170–1189. 10.1002/glia.22990.

39. Bernardos, R.L., and Raymond, P.A. (2006). GFAP transgenic zebrafish. Gene Expression Patterns 6, 1007–1013. 10.1016/j.modgep.2006.04.006.

40. Strata, P., and Harvey, R. (1999). Dale’s principle. Brain Res Bull 50, 349–350. 10.1016/S0361-9230(99)00100-8.

41. Eccles, J.C., Fatt, P., and Koketsu, K. (1954). Cholinergic and inhibitory synapses in a pathway from motor-axon collaterals to motoneurones. J Physiol 126, 524–562. 10.1113/jphysiol1954.sp005226.

42. Schlegel, P., Yin, Y., Bates, A.S., Dorkenwald, S., Eichler, K., Brooks, P., Han, D.S., Gkantia, M., dos Santos, M., Munnelly, E.J., et al. (2024). Whole-brain annotation and multi-connectome cell typing of Drosophila. Nature 634, 139–152. 10.1038/s41586-024-07686-5.

43. Shiu, P.K., Sterne, G.R., Spiller, N., Franconville, R., Sandoval, A., Zhou, J., Simha, N., Kang, C.H., Yu, S., Kim, J.S., et al. (2024). A Drosophila computational brain model reveals sensorimotor processing. Nature 634, 210–219. 10.1038/s41586-024-07763-9.

44. Brandt, J.P., and Smith, C.J. (2023). Piezo1-mediated spontaneous calcium transients in satellite glia impact dorsal root ganglia development. PLoS Biol 21, e3002319. 10.1371/JOURNAL.PBIO.3002319.

45. Green, L.A., Gallant, R.M., Brandt, J.P., Nichols, E.L., and Smith, C.J. (2022). A Subset of Oligodendrocyte Lineage Cells Interact With the Developing Dorsal Root Entry Zone During Its Genesis. Front Cell Neurosci 16, 1–20. 10.3389/fncel.2022.893629.

46. Green, L.A., Nebiolo, J.C., and Smith, C.J. (2019). Microglia exit the CNS in spinal root avulsion. PLoS Biol 17, 1–30. 10.1371/journal.pbio.3000159.

47. Nichols, E.L., Green, L.A., and Smith, C.J. (2018). Ensheathing cells utilize dynamic tiling of neuronal somas in development and injury as early as neuronal differentiation. Neural Dev 13, 19. 10.1186/s13064-018-0115-8.

48. Chalfie, M. (2009). Neurosensory mechanotransduction. Nat Rev Mol Cell Biol 10, 44–52.

49. Ernstrom, G.G., and Chalfie, M. (2002). Genetics of Sensory Mechanotransduction. Annu Rev Genet 36, 411–453. 10.1146/annurev.genet.36.061802.101708.

50. Way, J.C., and Chalfie, M. (1988). mec-3, a homeobox-containing gene that specifies differentiation of the touch receptor neurons in C. elegans. Cell 54, 5–16. 10.1016/0092-8674(88)90174-2.

51. Albeg, A., Smith, C.J., Chatzigeorgiou, M., Feitelson, D.G., Hall, D.H., Schafer, W.R., Miller, D.M., and Treinin, M. (2011). C. elegans multi-dendritic sensory neurons: Morphology and function. Molecular and Cellular Neuroscience 46, 308–317. 10.1016/j.mcn.2010.10.001.

52. Smith, C.J., Watson, J.D., Spencer, W.C., O’Brien, T., Cha, B., Albeg, A., Treinin, M., and Miller, D.M. (2010). Time-lapse imaging and cell-specific expression profiling reveal dynamic branching and molecular determinants of a multi-dendritic nociceptor in C. elegans. Dev Biol 345, 18–33. 10.1016/j.ydbio.2010.05.502.

53. Walker, D.S., Chew, Y.L., and Schafer, W.R. (2024). Genetics of C. elegans Behavior. In Oxford Research Encyclopedia of Neuroscience (Oxford University Press). 10.1093/acrefore/9780190264086.013.502.

54. Smith, C.J., O’Brien, T., Chatzigeorgiou, M., Clay Spencer, W., Feingold-Link, E., Husson, S.J., Hori, S., Mitani, S., Gottschalk, A., Schafer, W.R., et al. (2013). Sensory neuron fates are distinguished by a transcriptional switch that regulates dendrite branch stabilization. Neuron 79, 266–280. 10.1016/j.neuron.2013.05.009.

55. Li, W., Kang, L., Piggott, B.J., Feng, Z., and Xu, X.Z.S. (2011). The neural circuits and sensory channels mediating harsh touch sensation in Caenorhabditis elegans. Nat Commun 2, 315. 10.1038/ncomms1308.

56. Chalfie, M., and Sulston, J. (1981). Developmental genetics of the mechanosensory neurons of Caenorhabditis elegans. Dev Biol 82, 358–370. 10.1016/0012-1606(81)90459-0.

57. Abraira, V.E., and Ginty, D.D. (2013). The sensory neurons of touch. Neuron 79, 618–639. 10.1016/j.neuron.2013.07.051.

58. Watanabe, M., Dymecki, S.M., Chirila, A.M., Springel, M.W., Toliver, A.A., Zimmerman, A.L., Orefice, L.L., Bai, L., Song, B.J., Bashista, K.A., et al. (2017). The Cellular and Synaptic Architecture of the Mechanosensory Dorsal Horn. Cell. 10.1016/j.cell.2016.12.010.

59. Li, L., Rutlin, M., Abraira, V.E., Cassidy, C., Kus, L., Gong, S., Jankowski, M.P., Luo, W., Heintz, N., Koerber, H.R., et al. (2011). The functional organization of cutaneous low-threshold mechanosensory neurons. Cell 147, 1615–1627. 10.1016/j.cell.2011.11.027.

60. Lumpkin, E.A., and Caterina, M.J. (2007). Mechanisms of sensory transduction in the skin. Nature 445, 858–865. 10.1038/nature05662.

61. Bautista, D.M., and Lumpkin, E.A. (2011). Probing mammalian touch transduction. J Gen Physiol 138, 291–301. 10.1085/jgp.201110637.

62. Lumpkin, E.A., Marshall, K.L., and Nelson, A.M. (2010). The cell biology of touch. Journal of Cell Biology 191, 237–248. 10.1083/jcb.201006074.

63. Allen, N.J., and Lyons, D.A. (2018). Glia as architects of central nervous system formation and function. 185, 181–185.

64. Hardaway, J.A., Sturgeon, S.M., Snarrenberg, C.L., Li, Z., Xu, X.Z.S., Bermingham, D.P., Odiase, P., Spencer, W.C., Miller, D.M., Carvelli, L., et al. (2015). Glial Expression of the Caenorhabditis elegans Gene swip-10 Supports Glutamate Dependent Control of Extrasynaptic Dopamine Signaling. Journal of Neuroscience 35, 9409–9423. 10.1523/JNEUROSCI.0800-15.2015.

65. Ackerman, S.D., Perez-Catalan, N.A., Freeman, M.R., and Doe, C.Q. (2021). Astrocytes close a motor circuit critical period. Nature 592, 414–420. 10.1038/s41586-021-03441-2.

66. Ma, Z., and Freeman, M.R. (2020). Trpml-mediated astrocyte microdomain ca2+ transients regulate astrocyte-tracheal interactions. Elife 9, 1–18. 10.7554/ELIFE.58952.

67. Ziegenfuss, J.S., Doherty, J., and Freeman, M.R. (2012). Distinct molecular pathways mediate glial activation and engulfment of axonal debris after axotomy. Nat Neurosci 15, 979–987. 10.1038/nn.3135.

68. Ma, Z., Stork, T., Bergles, D.E., and Freeman, M.R. (2016). Neuromodulators signal through astrocytes to alter neural circuit activity and behaviour. Nature 539, 428–432. 10.1038/nature20145.

69. Kikuchi, T., Gonzalez-Soriano, J., Kastanauskaite, A., Benavides-Piccione, R., Merchan-Perez, A., DeFelipe, J., and Blazquez-Llorca, L. (2020). Volume Electron Microscopy Study of the Relationship Between Synapses and Astrocytes in the Developing Rat Somatosensory Cortex. Cerebral Cortex 30, 3800–3819. 10.1093/cercor/bhz343.

70. Murat, C.D.B., and García-Cáceres, C. (2021). Astrocyte Gliotransmission in the Regulation of Systemic Metabolism. Metabolites 11. 10.3390/metabo11110732.

71. Linne, M.-L. (2024). Computational modeling of neuron–glia signaling interactions to unravel cellular and neural circuit functioning. Curr Opin Neurobiol 85, 102838. 10.1016/j.conb.2023.102838.

72. Bazargani, N., and Attwell, D. (2016). Astrocyte calcium signaling: the third wave. Nat Neurosci 19, 182–189. 10.1038/nn.4201.

73. Kellner, V., Kersbergen, C.J., Li, S., Babola, T.A., Saher, G., and Bergles, D.E. (2021). Dual metabotropic glutamate receptor signaling enables coordination of astrocyte and neuron activity in developing sensory domains. Neuron 109, 2545-2555.e7. 10.1016/j.neuron.2021.06.010.

74. de Ceglia, R., Ledonne, A., Litvin, D.G., Lind, B.L., Carriero, G., Latagliata, E.C., Bindocci, E., Di Castro, M.A., Savtchouk, I., Vitali, I., et al. (2023). Specialized astrocytes mediate glutamatergic gliotransmission in the CNS. Nature 622, 120–129. 10.1038/s41586-023-06502-w.

75. Lu, T.-Y., Hanumaihgari, P., Hsu, E.T., Agarwal, A., Kawaguchi, R., Calabresi, P.A., and Bergles, D.E. (2023). Norepinephrine modulates calcium dynamics in cortical oligodendrocyte precursor cells promoting proliferation during arousal in mice. Nat Neurosci 26, 1739–1750. 10.1038/s41593-023-01426-0.

76. Duque, M., Chen, A.B., Hsu, E., Narayan, S., Rymbek, A., Begum, S., Saher, G., Cohen, A.E., Olson, D.E., Li, Y., et al. (2024). Ketamine induces plasticity in a norepinephrine-astroglial circuit to promote behavioral perseverance. Neuron. 10.1016/j.neuron.2024.11.011.

77. Thiele, T.R., Donovan, J.C., and Baier, H. (2014). Descending Control of Swim Posture by a Midbrain Nucleus in Zebrafish. Neuron 83, 679–691. 10.1016/j.neuron.2014.04.018.

78. Nichols, E.L., and Smith, C.J. (2019). Synaptic-like Vesicles Facilitate Pioneer Axon Invasion. Current Biology, 1–13. 10.1016/j.cub.2019.06.078.

79. Kucenas, S., Wang, W.-D., Knapik, E.W., and Appel, B. (2009). A Selective Glial Barrier at Motor Axon Exit Points Prevents Oligodendrocyte Migration from the Spinal Cord. Journal of Neuroscience 29, 15187–15194. 10.1523/JNEUROSCI.4193-09.2009.

80. Kimmel, C., Ballard, W., Kimmel, S., Ullmann, B., and Schilling, T. (1995). Stages of Embryonic Development of the Zebrafish. Developmental Dynamics 203, 253–310. 10.1002/aja.1002030302.

